# Enhancement of colorectal cancer therapy through interruption of the HSF1-HSP90 axis by p53 activation or cell cycle inhibition

**DOI:** 10.1101/2024.02.22.581507

**Authors:** Tamara Isermann, Kim Lucia Schneider, Florian Wegwitz, Tiago De Oliveira, Lena-Christin Conradi, Valery Volk, Friedrich Feuerhake, Björn Papke, Sebastian Stintzing, Bettina Mundt, Florian Kühnel, Ute M. Moll, Ramona Schulz-Heddergott

**Affiliations:** Department of Molecular Oncology, University Medical Center Göttingen, Göttingen, Germany; Charité – Universitätsmedizin Berlin, Institute of Pathology, Laboratory of Molecular Tumor Pathology and Systems Biology, Berlin, Germany; German Cancer Consortium (DKTK); Partner Site Berlin, German Cancer Research Center (DKFZ), Heidelberg, Germany; Department of Gynecology and Obstetrics, University Medical Center Göttingen, Göttingen, Germany; Department of General, Visceral, and Pediatric Surgery, University Medical Center Göttingen, Germany; Institute for Pathology, Hannover Medical School, Hannover, Germany; Charité – Universitätsmedizin Berlin, Department of Hematology, Oncology, and Cancer Immunology, Berlin, Germany; Department of Gastroenterology, Hepatology, Infectious Diseases and Endocrinology, Hannover Medical School, Hannover, Germany; Department of Pathology, Stony Brook University, Stony Brook, NY

**Keywords:** RG-7388, mutp53, p53, HSP90, cell cycle, CDK4/6, Palbociclib

## Abstract

The stress-associated molecular chaperone system is an actionable target in cancer therapies. It is ubiquitously upregulated in cancer tissues and enables tumorigenicity by stabilizing hundreds of oncoproteins and disturbing the stoichiometry of protein complexes. Most inhibitors target the key component heat-shock protein 90 (HSP90). However, although classical HSP90 inhibitors are highly tumor-selective, they fail in phase 3 clinical oncology trials. These failures are at least partly due to an interference with a negative feedback loop by HSP90 inhibition, known as heat-shock response (HSR): in response to HSP90 inhibition there is compensatory synthesis of stress-inducible chaperones, mediated by the transcription factor heat-shock factor 1 (HSF1). We recently identified that wildtype p53 (p53) actively reduces the HSR by repressing HSF1 via a p21-CDK4/6-MAPK-HSF1 axis. Here we test the hypothesis that in HSP90-based therapies simultaneous p53 activation or direct cell cycle inhibition interrupts the deleterious HSF1-HSR axis and improves the efficiency of HSP90 inhibitors.

Indeed, we find that the clinically relevant p53 activator Idasanutlin suppresses the HSF1-HSR activity in HSP90 inhibitor-based therapies. This combination synergistically reduces cell viability and accelerates cell death in p53-proficient colorectal cancer (CRC) cells, murine tumor-derived organoids and patient-derived organoids (PDOs). Mechanistically, upon combination therapy human CRC cells strongly upregulate p53-associated pathways, apoptosis, and inflammatory immune pathways. Likewise, in the chemical AOM/DSS CRC model in mice, dual HSF1-HSP90 inhibition strongly represses tumor growth and remodels immune cell composition, yet displays only minor toxicities in mice and normal mucosa-derived organoids. Importantly, inhibition of the cyclin dependent kinases 4 and 6 (CDK4/6) under HSP90 inhibition phenocopies synergistic repression of the HSR in p53-proficient CRC cells. Even more important, in p53-deficient (mutp53-harboring) CRC cells, an HSP90 inhibition in combination with CDK4/6 inhibitors similarly suppresses the HSF1-HSR system and reduces cancer growth. Likewise, p53-mutated PDOs strongly respond to dual HSF1-HSP90 pathway inhibition and thus, providing a strategy to target CRC independent of the p53 status.

In sum, activating p53 (in p53-proficient cancer cells) or inhibiting CDK4/6 (independent of the p53 status) provide new options to improve the clinical outcome of HSP90-based therapies and to enhance colorectal cancer therapy.

## INTRODUCTION

Colorectal cancer (CRC) is the second most common cause of cancer-related deaths in industrialized countries leading to nearly 1 million deaths per year [1]. In 2020, CRC was estimated to count for 10% of all new cancer diagnoses and 9.4% of all cancer deaths [2]. Advanced and metastasized CRC stages have a 5-years survival rate of approximately 15% [Morris ASCO]. Thus, new therapeutic strategies are urgently needed, especially for advanced stages.

A promising target in cancer therapy is the chaperone system. During oncogenesis the normal chaperone function (aiding other proteins in proper folding) is dramatically subverted to enable the malignant lifestyle of tumor cells [3–11]. Consequently, cancer cells are strongly addicted to a ubiquitously upregulated chaperone system [8, 12–14]. The main reason is that cancer cells are under constant and cumulative proteotoxic stress, driven by most hallmarks of cancer [15]. They defend against these pathophysiological stresses by constitutive activation of heat-shock factor 1 (HSF1), the master transcription factor of stress-induced chaperones [16–18]. Thus, HSF1 hyper-activation leads to massive up-regulation of stress-inducible heat-shock proteins (HSPs), foremost HSP90α, plus HSP70, other HSPs and numerous pro-tumorigenic co-chaperones. This cumulates in the formation of tumor cell-specific super-chaperone complexes with HSP90 as key component. Consequently, HSP90 stabilizes hundreds of mutated and truncated oncoproteins which normally would be degraded by proteasomes. HSP90-stabilized oncogenic proteins, collectively called HSP90 clients, include missense p53 mutants (mutp53) [11, 19–22] and Bcr-Abl fusions [20], overexpressed receptor tyrosine kinases (ErbB1, ErbB2/HER2) [16, 23], signaling kinases (AKT) [24], hormone receptors and cytokines [16, 25].

Since cancer cells are addicted to the HSF1-regulated chaperone system, it represents an attractive tumor-selective therapeutic target. Most of the promising small molecule inhibitors were generated against the N-terminal ATPase domain of HSP90 [6, 7, 26–28]. However, despite encouraging pre-clinical results, most if not all clinical oncology trials using HSP90 inhibitors alone or in combination with chemotherapeutics have failed so far [6, 7, 26, 28]. We hypothesized that a major reason for the failure might be the compensatory stimulation of HSF1 by the inadvertent interference of the HSF1-HSP90 negative feedback loop upon HSP90 inhibition. This severely limits the therapeutic efficacy of HSP90 inhibitors. In normal cells, this feedback via inhibitory HSP90-HSF1 protein-protein binding is needed to properly control HSF1 activity. Normally, upon activation HSF1 trimerizes and binds to its cognate DNA element to induce stress-inducible HSPs such as HSP90α, HSP70s, HSP40s and co-chaperones – a reaction known as the heat-shock response (HSR) [3–10]. However, to limit and control HSF1 activation, the target protein HSP90 - through its N-terminal domain - binds back and inactivates HSF1, thus establishing a negative feedback loop. In contrast, upon proteotoxic stress HSP90 dissociates from HSF1 to re-activate the HSR. Since cancer cells are under constant proteotoxic stress, their negative feedback loop is greatly diminished and HSF1 is partially reactivated as a chronic condition. Pharmacological HSP90 inhibition phenocopies HSP90 disappearance and thus promotes binding and activation of HSF1 to its cognate DNA response element, thereby fully inducing the unwanted HSR in cancer cells. As a result, tumor-driving proteins might be further stabilized by alternative chaperones other than HSP90 [29–32]. Thus, preventing the deleterious HSR upon HSP90 inhibition in cancer cells is predicted to improve the therapeutic efficacy of clinically relevant HSP90 inhibitors.

Our previous findings might provide a viable approach towards this goal. We recently showed that wildtype p53 (here termed ‘p53’) suppresses HSF1 activity in human CRC cells, tumor-derived organoids and in a CRC model in mice *in vivo* [33]. p53, a key tumor suppressor preventing tumor initiation and tumor progression, is activated by a broad spectrum of stresses such as DNA damage, oncogene hyperactivation, hypoxia or metabolic stress [20, 34, 35]. In response, p53 transcriptionally induces genes to maintain cell integrity through e.g. transient cell cycle arrest, activation of DNA repair, and, if cells are irreparably damaged via induction of cell death. We found that activated p53 suppresses HSF1 activity and its target genes (i.e. the HSR) via the downstream CDKN1A/p21 - CDK4/6 - E2F - MLK3 – MAPK axis [33]. Specifically, the p53 - p21 axis induces a cell cycle arrest via CDK4/6 inhibition. Based on these findings, we asked whether HSF1 suppression by either p53 activation or CDK4/6 inhibition prevents the HSR in the context of HSP90 inhibition, thereby allowing a far more profound suppression of chaperone activity. Indeed, p53 activation or CDK4/6 inhibition strongly antagonizes the unwanted activation of HSF1 and HSR upon treatment with HSP90 inhibitors. In addition to HSP90 inhibition we show here that dual pathway inhibition by p53 activation via RG-7388 (Idasanutlin, a clinically relevant p53 activator) prevents upregulation of HSF1 target gene expression in human CRC cells. Importantly, combination treatment synergistically reduces cell viability and increases apoptosis in CRC cell lines, murine CRC-derived organoids and in patient-derived organoids (PDOs). Notably, in combination therapy, inflammatory pathways are strongly upregulated, indicated by deregulation of immune cell composition in our CRC mouse model *in vivo*. Dual pathway inhibition profoundly diminishes tumor sizes without obvious toxicities for mice. Likewise, in p53-deficient CRC cells and PDOs, CDK4/6 inhibition by Palbociclib phenocopies p53 activation and lowers the HSR by repressing HSF1 target genes, causing reduced cancer cell growth.

In sum, our results show that p53 activation or CDK4/6 inhibition can each prevent the HSR rebound response induced by HSP90 inhibition in tumor cells. Dual inhibition improves HSP90-based therapies by *i)* abrogating the HSR under HSP90 inhibition, *ii)* more efficient degradation of HSP90 clients and, *iii)* additional positive effects from the p53 tumor suppressive program or from cell cycle arrest. This improves the therapeutic effect of the promising class of highly tumor-selective HSP90 inhibitors, independent of the p53 status and, independent of the first-line CRC therapy status.

## RESULTS

### Dual HSF1-HSP90 inhibition via p53 activation synergistically impairs colorectal cancer cell growth by abrogating the heat-shock response

HSP90 inhibitors (HSP90i) are highly tumor selective and represent a promising class of anti-cancer drugs [3–11, 16, 20, 25, 36–39]. Although HSP90 inhibition alone shows strong anti-tumorigenic effects in experimental mouse models in vivo, most HSP90 inhibitors have so far failed in clinical trials, one reason being the rebound HSF1-induced heat-shock response (HSR) upon HSP90 inhibition [5, 10, 13, 26]. Since we recently demonstrated that the HSF1-governed HSR is suppressed by wildtype p53 (p53) activation upon Nutlin-3a [33], we tested the strategy of counter-regulating the unwanted HSR through HSF1 suppression by p53 activation in p53-proficient tumors upon HSP90-based therapies. Approx. 40% of human CRC tumors harbor wildtype p53 presenting a significant proportion of colorectal cancer (CRC) patients.

To activate p53, we used the clinically advanced Nutlin-3a derivate Idasanutlin (RG-7388), an MDM2 inhibitor currently in numerous clinical trials. MDM2 is the principal antagonist of p53. Idasanutlin disrupts p53’s interaction with MDM2 and thus blocks its E3 ligase-mediated proteasomal degradation, thereby stabilizing and activating p53 [35, 40, 41]. We chose this highly specific non-genotoxic mode of p53 activation to avoid generating confounding DNA damage with its multiple activated pathways that can contribute to mutually contradictory HSF1 modifications [42, 43]. For HSP90 inhibition we used Ganetespib and Onalespib, both currently also in clinical trials.

Indeed, as shown in Figs. 1A and S1A, combination treatment of Ganetespib plus RG-7388 yielded additive to synergistic reduction of cell viability in p53-proficient CRC cell lines HCT116 and RKO, compared to single treatments. Combination treatments showed stronger cell viability effects for all tested drug concentrations compared to corresponding single drugs of the same concentration (Figs. 1A and S1A). Of note, synergistic effects were p53-dependent. In contrast to p53-proficient cells, isogenic HCT116 cells harboring a homozygous p53 deletion failed to show stronger effects upon dual HSF1-HSP90 pathway inhibition (Fig. S1B).

**Figure 1.**
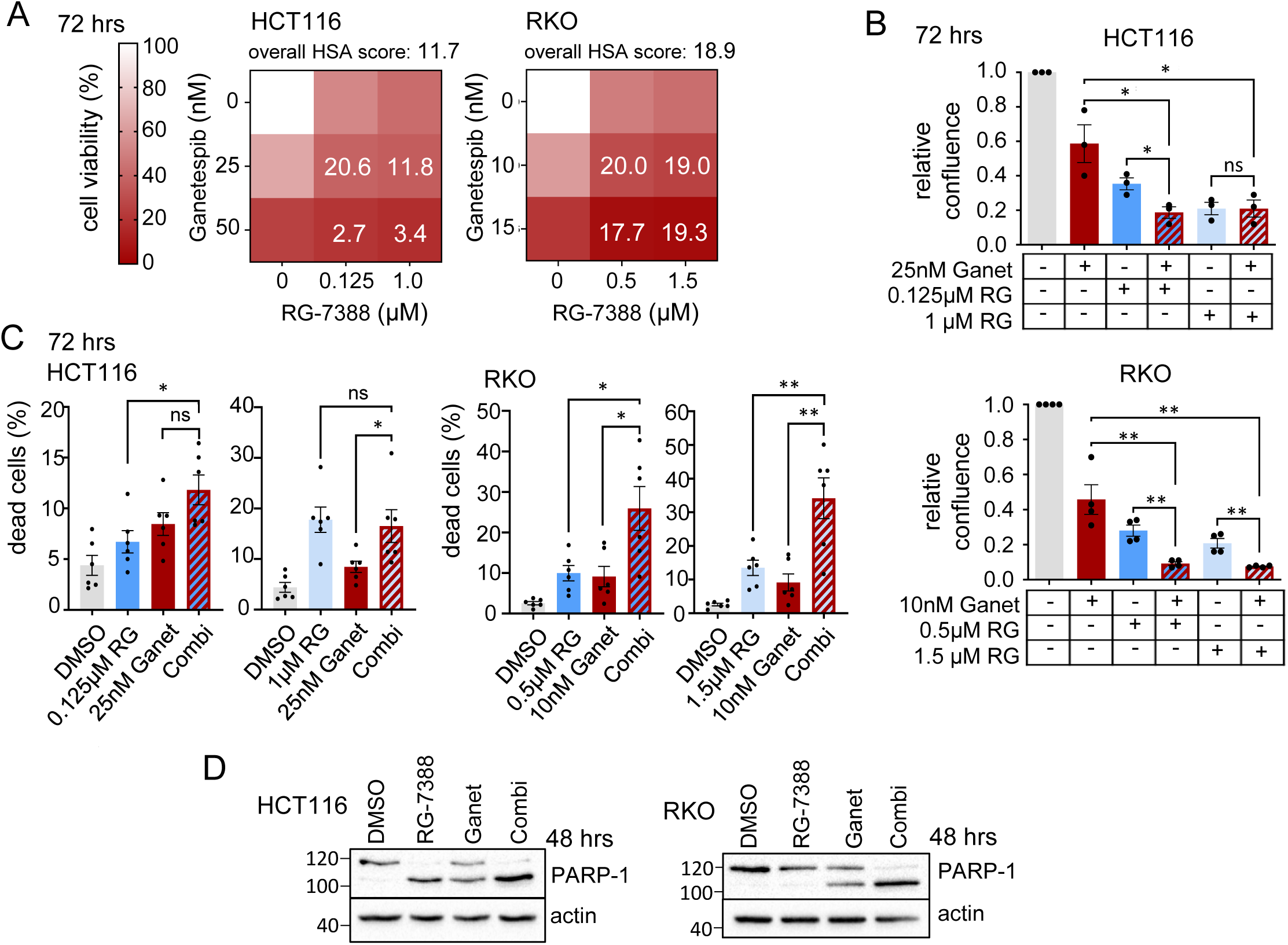
Dual HSP90-HSF1 inhibition via p53 activation synergistically impairs colorectal cancer cell growth. (A) Cell viability matrices of HCT116 (left) and RKO (right) cells treated with Ganetespib – RG-7388 combination for 72 hrs at indicated concentrations. Color scheme represents changes in cell viability. Numbers within the matrix indicate the HSA synergy score. Synergy scores: < −10 antagonistic; −10 to 10 additive; > 10 synergistic. ≥ 3 biological replicates. (B) Relative confluence after 72 hrs treatment of HCT116 (top) and RKO (bottom) cells. Cell confluence was analyzed by Celigo imaging cytometer. Confluence relative to DMSO control, set at value 1. (C) Induction of cell death. PI/Hoechst/Annexin V staining of HCT116 (left) and RKO (right) cells treated for 72 hrs with Ganetespib and RG-7388 at the indicated concentrations. Percent dead cells include PI+ only, annexin V+ only and PI+Annexin V+ cells and were analyzed by Celigo imaging cytometer. (D) PARP-1 immunoblots of HCT116 (left) and RKO (right) cells treated for 48 hrs. HCT116 were treated with 50nM Ganet and 1μM RG. RKO were treated with 25nM Ganet and 1.5μM RG. Representative immunoblots shown from 3 replicates with 2 independent experiments each. (B-C) Mean ± SEM from ≥ 3 biological replicates, Student’s t-test, p*≤ 0.05, p**≤ 0.01, p***≤ 0.001; ns, not significant. Ganet: Ganetespib, RG: RG-7388.

Similarly, cancer cell confluence decreased further upon p53 activation with combinatorial HSP90i plus RG-7388 treatment, compared to single treatments (Figs. 1B and S1C). Likewise, combination treatments induced stronger cell death at different drug concentrations, compared to single treatments (Fig. 1C). Although cell death was only modest in HCT116 cells (Fig. 1C), PARP cleavage was readily detectable in both cell lines (Fig. 1D). In line, a shorter 48 hrs dual pathway inhibition also increased cell death in RKO cells, but not in HCT116 cells compared to single treatment (Fig. S1D). p53 activation by RG-7388 also diminished cell viability upon combinatorial Onalespib treatment compared to single treatments (Fig. S1E), although the effects were milder than for Ganetespib (compare to Fig S1A).

Since we hypothesized that the observed synergistic effects arise from the abrogation of the HSP90-HSF1 negative feedback loop, we analyzed the heat shock response (HSR) upon concomitant p53 activation in HSP90-based therapy, using quantitative real-time PCR. Indeed, we found that dual HSF1-HSP90 pathway inhibition suppresses the increase of HSF1 target gene expression compared to Ganetespib alone in HCT116 and RKO cells, using a range of drug concentrations (Figs. 2A and S2A). Randomly selected classical HSF1 targets including *HSPH1*, *HSPE1*, *HSPB1*, *HSPA1A* and *HSP90AB1* were downregulated by concomitant p53 activation during HSP90 inhibition. While not every HSF1 target gene is upregulated upon Ganetespib alone (Figs. 2A and S2A, HSR response, compare grey with red bars), a dual HSF1-HSP90 pathway inhibition mostly repressed them below the level of untreated control cells (DMSO), reflecting a diminished basal HSF1 activity. Moreover, RNAseq analysis confirmed that p53 activation combined with HSP90 inhibition reduces global HSF1 target gene expression, compared to the HSR-inducing Ganetespib single treatment (Fig. 2B). The critical phosphorylation site for HSF1 activation is on Ser326 (pHSF1) which serves as functional hallmark for the tumor-promoting HSR [12, 16]. Consistent with the observed HSF1 target gene repression, RG-7388 in combination with Ganetespib strongly reduced pSer326-HSF1 levels in HCT116 and RKO cells compared to HSP90 inhibition alone (Fig. 2C). Moreover, RG-7388 combined with Ganetespib destabilizes classical HSP90 clients such as AKT and cRAF more strongly than Ganetespib alone (Fig. 2D), confirming the pronounced inactivation of the HSF1 - HSR response and consequently, profound destabilization of HSP90 clients. To further understand the consequences of a dual HSF1-HSP90 pathway repression, RNAseq data were subjected to GSEA pathway analysis (Figs. 2E, F). Indeed, compared to RG-7388 treatment alone (grey bars), RG-7388 in combination with Ganetespib broadly enhanced the hallmarks of p53-associated pathways (Fig. 2E, red bar). The same is seen when comparing the combination versus Ganetespib alone (Fig. 2F, red bar). Conversely, compared to RG-7388 or Ganetespib alone, combination treatment impaired cell cycle-driving pathways including E2F target genes, MYC pathways and the G2M checkpoint more strongly (Figs. 2E, F).

**Figure 2.**
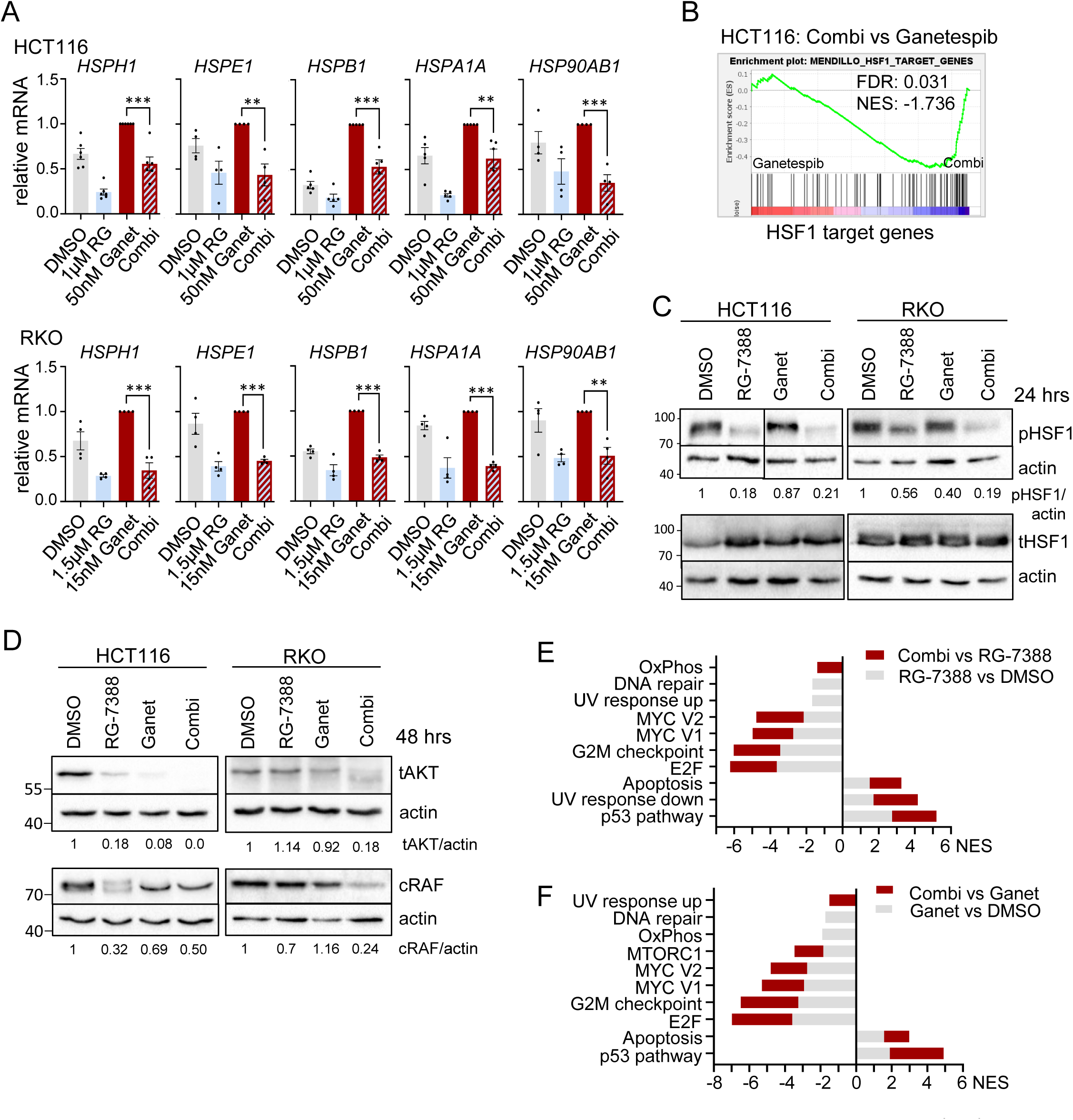
Dual HSP90-HSF1 inhibition via p53 activation abrogates the negative feedback loop and destabilizes HSP90 clients. (A) mRNA expression levels of HSF1 target genes in HCT116 (top) and RKO (bottom) cells treated for 24 hrs. qRT-PCRs for the indicated mRNAs normalized to *RPLP0* mRNA. Mean ± SEM from ≥ 3 biological replicates. Student’s t-test, p*≤ 0.05, p**≤ 0.01, p***≤ 0.001; ns, not significant. (B) GSEA enrichment plot for HSF1 target genes in HCT116 cells treated with drug combination (1µM RG-7388 and 50nM Ganetespib) versus Ganetespib-only (50 nM). HSF1 target gene list from Mendillo *et al.* [62]. (C, D) Immunoblots for HCT116 and RKO cells treated with 1µM RG-7388, 50nM Ganetespib for HCT116, or 1.5µM RG-7388, 15nM Ganetespib for RKO cells, treated for 24 hrs (C) or for 48 hrs (D). (C) Blots were probed for phospho-Ser326 HSF1 (pHSF1) and total HSF1 (tHSF1). (D) Blots were probed for total AKT (tAKT) and cRAF. Densitometric expression levels normalized for actin are indicated. (E, F) GSEA pathway analysis for hallmark gene sets on RNAseq data from HCT116 cells treated for 24 hrs as in (B), comparing drug combination versus RG-7388 alone (E) or Ganetespib alone (F). Grey bars, NES (Normalized Enrichment Score) of RG-7388 (E) or Ganetespib (F) treatment relative to DMSO. Red bars, NES of combination treatment relative to either RG-7388 or Ganetespib treatment alone, respectively.

In sum, we conclude that p53 activation effectively suppresses the HSF1-mediated HSR and synergistically impairs human CRC cell growth, thereby improving HSP90-based therapies. Moreover, the powerful tumor-suppressive p53 program can be profoundly activated in the presence of concomitant HSP90 inhibition.

### Activated p53 represses the heat-shock response in HSP90-based therapies in murine colorectal tumor-derived organoids and CRC patient-derived organoids

To further strengthen the evidence for p53-mediated improvement of HSP90-based therapies, we generated colorectal tumor-derived organoids from the AOM/DSS mouse model. To this end, we co-treated wildtype p53 organoids with RG-7388 plus Ganetespib and observed dampened cell viability with additive to synergistic effects versus each drug alone (Fig. 3A). This was accompanied by suppression of HSF1 target genes upon different concentrations of Ganetespib (Fig. 3B). Importantly, cell death was minimal and did not significantly change in normal mucosa-derived organoids generated from the small intestine (Fig. 3C). In contrast, the same combined drug concentrations increased cell death impressively to up to 80% in CRC-derived organoids (Fig. 3D), underlining the tumor-selectivity and therapeutic value of dual HSF1-HSP90 pathway inhibition. Moreover, we generated patient-derived organoids (PDOs) from 2 different wtp53 CRC patients, one from a primary tumor (PDO #1) and one from a liver metastasis (PDO #2). Both PDO cultures confirmed the enhanced defect in cell viability with drug combinations compared to the corresponding single drugs (Figs. 3E, F). Since drug responses can be dependent on the organoid volume [44], we also tested different organoid sizes. Larger, multi-volume organoids (Fig. 3E, PDO #2) as well as single cell-dissociated smaller organoids (Fig. 3F) generated from the same PDO culture, both favorably responded to dual HSF1-HSP90 inhibition with stronger reduced viability and reduction of organoid sizes and numbers compared to single treatments.

**Figure 3.**
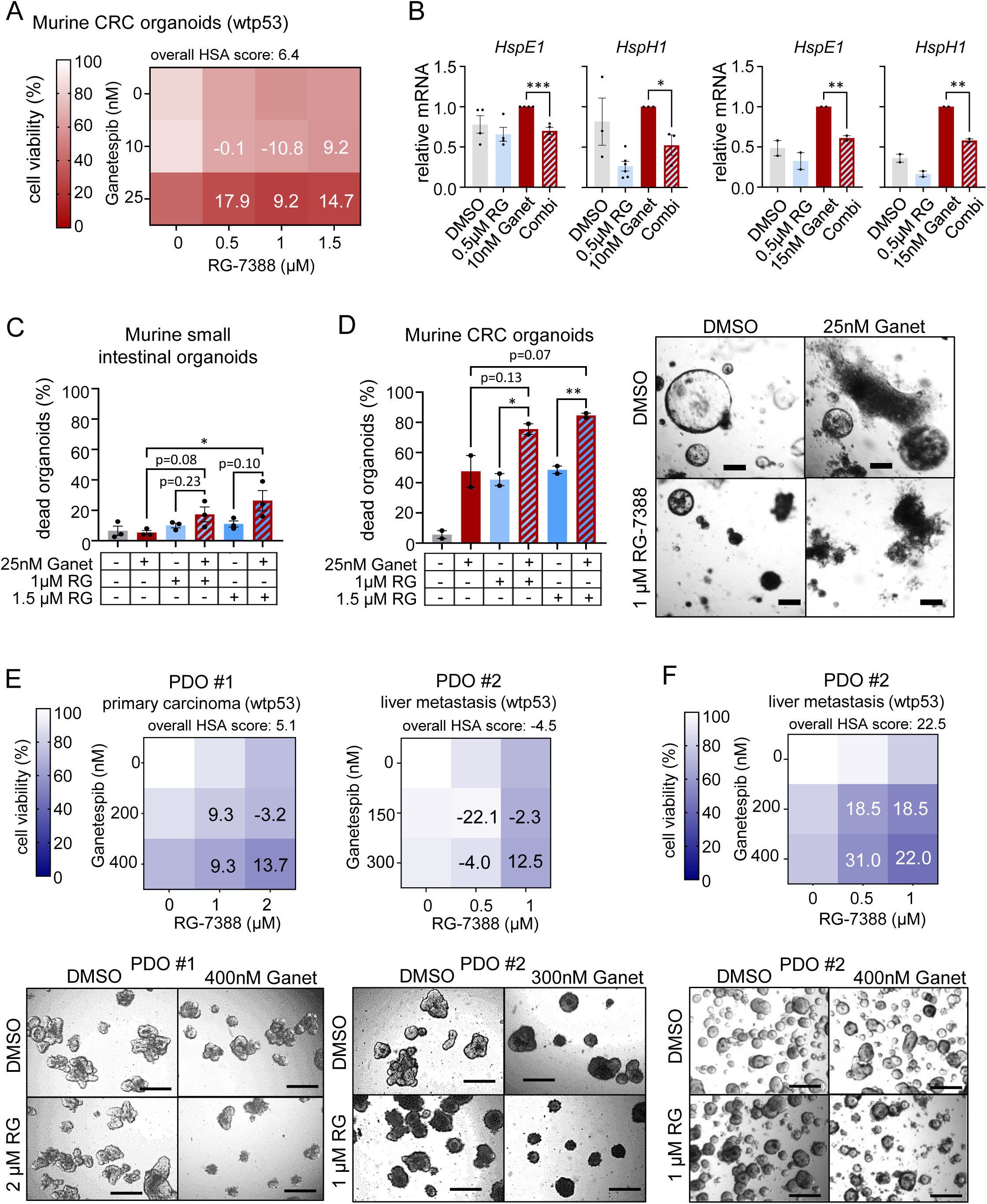
Activated p53 represses the HSF1-mediated heat-shock response in HSP90-based therapies in murine and human colorectal tumor-derived organoids. (A) Cell viability matrix of murine CRC tumor organoids treated with Ganetespib – RG-7388 combination treatment for 48 hrs. Organoids generated from AOM/DSS treated C57BL6/J mice. Four independent biological replicates with 3 in-plate technical replicates each. (B) mRNA expression levels of HSF1 target genes in murine CRC tumor organoids generated from AOM/DSS mice treated with 10nM (left) and 15nM (right) Ganetespib in combination with 500 nM RG-7388 for 24 hrs. qRT-PCRs for the indicated mRNAs normalized to *RPLP0* mRNA. Mean ± SEM from min. 2 independent biological replicates. Student’s t-test, p*≤ 0.05, p**≤ 0.01, p***≤ 0.001; ns, not significant. (C) Quantification of normal small intestinal mucosa-derived organoids treated with the indicated combinations for 48 hrs. Mean ± SEM from 3 independent biological replicates. Student’s t-test, p*≤ 0.05, p**≤ 0.01, p***≤ 0.001; when ns, p-value indicated. (D) *left*, Quantification of dead CRC tumor-derived organoids treated with drug combination for 48 hrs as in (C). Organoids were generated from AOM/DSS treated C57BL6/J mice. Mean ± SEM from 2 independent biological replicates. *right*, Representative brightfield images of CRC tumor organoids. Scale bars, 100 μm. Ganet: Ganetespib, RG: RG-7388. (E) Cell viability matrices of p53-proficient (wildtype p53, wtp53) patient-derived organoids (PDOs) treated with Ganetespib, RG-7388 or in combination for 48 or 72 hrs. A PDO case represents one PDO culture (PDO #). For each PDO culture, four replicates (different passages) with 3 in-plate technical replicates each were measured. Organoids were cultivated and treated as large organoids. *Bottom*, Representative brightfield images of large CRC tumor organoids. Scale bars, 200 μm. (F) Cell viability matrix of PDO #2 culture, cultivated and treated as small organoids with Ganetespib, RG-7388 or in combination for 3 or 5 days. Three replicates (different passages) with 3 in-plate technical replicates each were measured. *Bottom*, Representative brightfield images of small PDOs #2 culture. Scale bars, 200 μm. (A, E, F) Color scheme represents changes in cell viability. Numbers in the matrix are HSA synergy scores. Synergy scores: < −10 antagonistic; −10 to 10 additive; > 10 synergistic. Ganet: Ganetespib, RG: RG-7388.

### Dual HSF1-HSP90 pathway inhibition reduces tumor progression in a CRC mouse model by remodeling the immune cell composition

The induction of the heat shock response is a major problem associated with HSP90 inhibitors in preclinical and clinical trials [18, 45–48]. Due to the negative feedback loop between HSP90 and HSF1, inhibition of HSP90 alone inadvertently upregulates HSF1 and other HSPs like HSP40 and HSP70 members, thereby stabilizing client proteins and dampening the effect of HSP90 inhibition. Thus, we examined in the murine AOM/DSS-induced autochthonous CRC model whether the combination treatment would improve HSP90-based therapy *in vivo*. This CRC model mirrors human CRC at the pathological and molecular level by exclusively generating tumors in the colorectal part of the intestine [49].

Mice with a defined tumor burden, assessed and scored by regular colonoscopy, were treated with either RG-7388 and Ganetespib in combination or drug alone for 3 weeks as indicated (Fig. 4A). Indeed, p53 activation concomitant with HSP90 inhibition strongly reduced the sizes and numbers of established colonic tumors (Fig. 4B). Gross pathological and histological analysis confirmed that combo-treated mice compared to single-treatment mice showed the highest degree of tumor shrinkage and largest reduction in tumor numbers (consistent with tumor prevention) (Figs. 4B, C, D and S4A). Combination treatment decreased the tumor area by approx. 50 - 60% compared to monotherapies (Fig. 4D). Weight loss is a sensitive indicator of toxicity of cancer therapies. However, the combo-treated mice remained physically active and did not lose weight, indicating a favorable and tolerable toxicity profile in mice (Fig. S4B).

**Figure 4.**
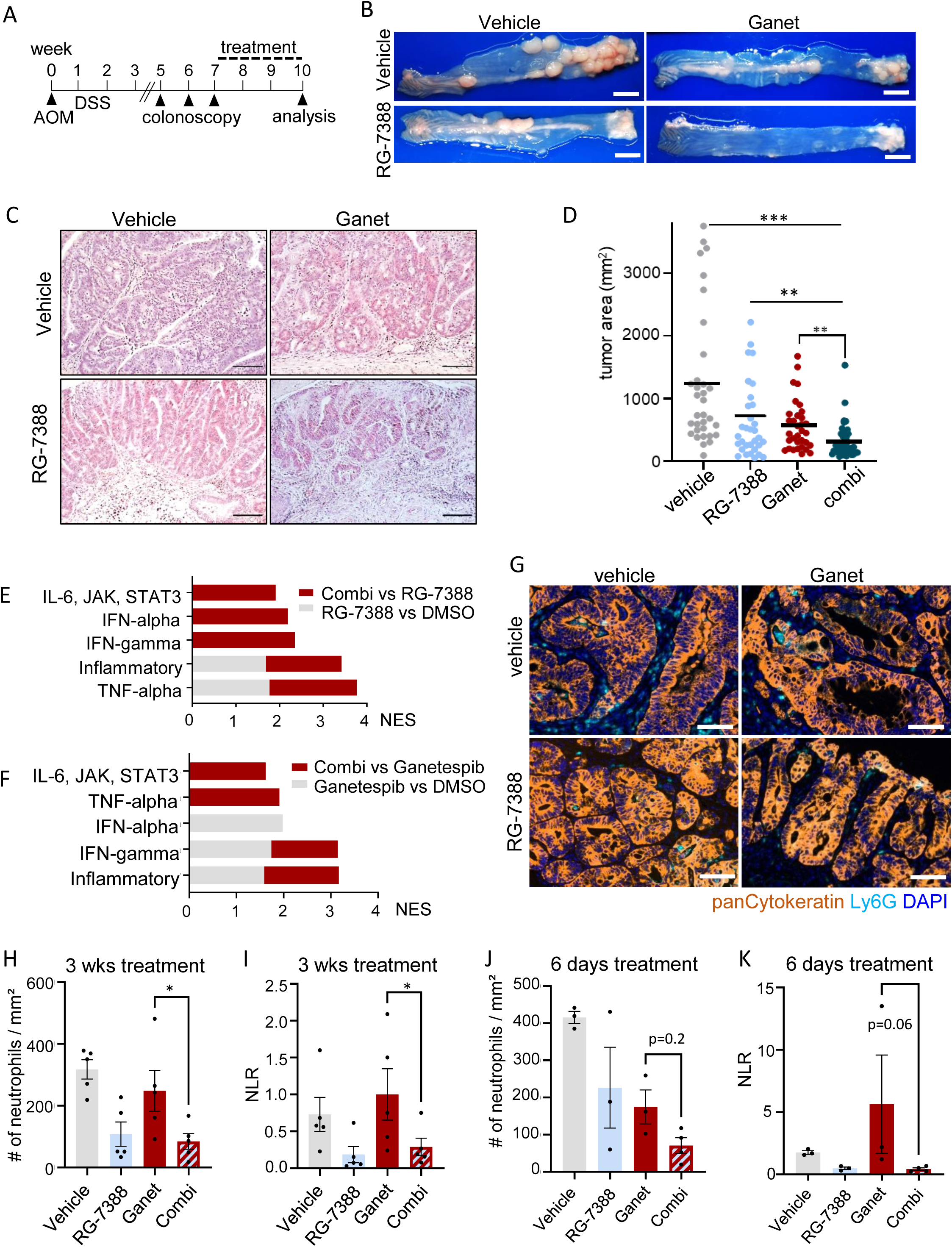
Dual HSF1-HSP90 pathway inhibition reduces tumor progression in a CRC mouse model by remodeling the immune system. (A) Treatment scheme for the p53-proficient AOM/DSS mouse model. After tumor visualization by repeat colonoscopy and tumor scoring for at least one S2-sized tumor and three S1-sized tumors per mouse, mice were treated with single drugs or combination treatment for 19 days. 4 hrs after the final dose, mice were dissected and analyzed. (B, C) Representative images of entire resected colons (B) and H&E-stained colon sections (C) from single or combination treated C57BL6/J mice at endpoint as described in (A). Mice received 50 mg/kg RG-7388 orally 5x per week, or 50 mg/kg Ganetespib intravenously 2x per week, or both. (B) scale bar 5 mm. (C) x20 magnification, scale bars, 100 µm. (D) Tumor surface area from AOM/DSS mice that had received single or combination treatment for 19 days as in (A). Cross-sections of swiss roles were H&E stained (Supp Fig. 4A). Tumor areas of all tumors per H&E-stained swiss role were measured using ImageJ and calculated as ellipsoid in mm^2^. Bar, mean. vehicle n = 35 tumors from 5 mice; RG-7388 and Ganetespib n = 33 tumors from 6 mice each; combination n = 45 tumors from 9 mice. (E, F) GSEA hallmark analysis of RNAseq data from HCT116 cells treated with drug combination or single drugs for 24 hrs. Enriched pathways are plotted according to NES. Grey bars: Enrichment in (F) RG-7388-only or (G) Ganetespib-only versus DMSO controls. Red bars: further enrichment with combination treatment relative to single treatment. NES: normalized enrichment score. (G-I) Multiplex immunohistochemistry of p53-proficient tumors from AOM/DSS mice receiving drug treatments for 19 days as in (A). (G) Representative images of indicated groups. Ly6G staining represents neutrophils and pan-Cytokeratin represents tumor epithelial cells. DAPI counter staining. Scale bars, 50 µm. (H) Quantification of Ly6G+CD11b+F4/80-neutrophils using the “Vectra Polaris” platform and the inForm Advanced Image Analysis software. n = 5 mice per group. All tumors from stained swiss roles were analyzed for indicated groups. (I) Plot of the neutrophil-lymphocyte-ratio (NLR) as marker of an inflammatory response. A low NLR indicates an inflammatory response. (J, K) Multiplex immunohistochemistry of tumor-bearing AOM/DSS mice who received a short drug treatment of 6 days. Quantification of Ly6G+CD11b+F4/80-neutrophils of the indicated groups as in (H). DMSO, Ganetespib and RG-7388 mice n = 3 each, combination treated mice n = 4. (C, H-K) Mean ± SEM. Student’s t-test, p*≤ 0.05, p**≤ 0.01, p***≤ 0.001; ns, not significant. Ganet: Ganetespib, RG: RG-7388

To mechanistically dissect the growth reduction upon dual HSF1-HSP90 inhibition, we further analyzed our RNAseq data (Figs. 2E, F). Of note, we observed massive upregulation of inflammatory hallmarks specifically with combinatorial treatment in human HCT116 cells (Figs. 4E, F). Inflammatory pathways including IL-6-JAK-STAT3, TNFα and Interferon signaling were either exclusively upregulated in response to the combination treatment (red bars) or were further increased in combination treatment compared to single RG-7388 (Fig. 4E) or Ganetespib treatment (Fig. 4F) (grey - red stacked bars). The GSEA data points to a modulation of immune cell regulatory pathways. Consistently, it is known that cancer-specific alteration of the p53-status affects immune activation and intratumoral composition of immune cell populations [50, 51]. Therefore, we performed a multiplex immunohistological (mIHC) analysis to investigate major lymphocytic and myeloid immune cell populations by staining with antibodies for CD4, CD8, Ly6G, CD11b, and F4/80 together with the tumor epithelial cell marker panCytokeratin in our CRC mouse model (Figs. 4A, G). The panCytokeratin staining in mIHC analysis mirrors the reduction of tumor mass after three weeks of treatment (Figs. 4A-D, Fig. 4G). CD4+ T cells, CD8+ T cells, and F4/80+ macrophages displayed no significant changes between treatment groups (data not shown). In contrast, Ly6G+CD11b+F4/80- cells were effectively decreased in combination treatment compared to HSP90-based inhibition (Fig. 4G, H). These cells, further shortly referred to as neutrophils, contain both tumor-associated neutrophils (TAN) and the myeloid-derived suppressor cell subset of neutrophils (PMN-MDSC). The protumorigenic and immunosuppressive role of neutrophils has been increasingly recognized [52]. Reflecting a predominantly myeloid and immunosuppressive character of the tumor microenvironment, a high ratio of neutrophils is an important predictor for a poor clinical outcome in many solid tumors, including CRC [53–58]. Consistent with the observed tumor reduction by our tumor-suppressive combination treatment, we assume that the combination treatment effectively reduced the immunosuppressive intratumoral neutrophil population suggesting an improved intratumoral immunoactivation. A decreased neutrophil-to-lymphocyte-ratio (NLR; Ly6G+ CD11b+ F4/80- to CD8+) in combo-treated mice compared to Ganetespib treated mice therefore reflects the conversion of the tumor microenvironment into a more immunoactivated state. Furthermore, short drug treatments in the AOM/DSS mouse model again decreased neutrophils (Fig. 4J) and NLR ratio (Fig. 4K) further strengthening that p53 activation in HSP90-based therapies effectively remodels the intratumoral immune cell composition and promotes immune activation.

In sum, we demonstrate a higher pre-clinical efficacy with dual HSF-HSP90 pathway inhibition in a CRC mouse model *in vivo* and confirm that in this synergistic context HSP90 inhibitors are promising cancer drugs with strong tumor selectivity.

### CDK4/6 inhibition in combination with HSP90 inhibitors impairs viability of p53-deficient cancer cells

The second most common alterations in CRC after APC mutations are bi-allelic *TP53* mutations (typically with LOH of the second allele), affecting over 60% of CRC patients [20] [https://www.cbioportal.org/]. The event of *TP53* mutations typically enables the critical colonic adenoma-to-carcinoma transition into frank invasive cancer and drives tumor aggressiveness [59–61]. Consequently, patients with p53-deficient CRC would not benefit from p53 activators in the context of HSP90 inhibition (see Fig. S1B, HCT116 p53-/- cells). Since we identified previously that p53 suppresses HSF1 via a repressive CDKN1A/p21 – CDK4/6 axis [33], we hypothesized that direct CDK4/6 inhibitors phenocopy the wtp53 – CDKN1A/p21 activation axis.

To this end we first tested whether CDK4/6 inhibition in general contributes to dual HSF1-HSP90 pathway inhibition, focusing on p53-proficient cells. Indeed, in HCT116 cells Palbociclib, a clinically used CDK4/6 inhibitor, caused marked reduction of cell viability in combination with Ganetespib (Figs. 5A, B) or Onalespib (Fig. 5C) in HSP90-based therapies. Cell confluence upon Palbociclib plus Ganetespib was strongly decreased, compared to Ganetespib alone (Figs. 5D and S5A). Furthermore, CDK4/6-associated dual pathway repression markedly increased cell death, as measured by PI/AnnexinV staining (Figs. 5E and S5B) and PARP cleavage by immunoblot (Fig. 5F), compared to single Ganetespib or Palbociclib treatment. Interestingly, in p53-proficient RKO cells additional CDK4/6 inhibition upon HSP90 inhibition showed only modest differences in cell viability (Figs. S5C-E), cell confluence (Fig. S5F, G) and cell death (Figs. S5H-J). Importantly however, in patient-derived organoids (PDOs), a clinically relevant translational model system generated from a CRC liver metastasis with wildtype p53, Palbociclib in combination with Ganetespib strongly diminished organoid viability and size (Fig. 5G).

**Figure 5:**
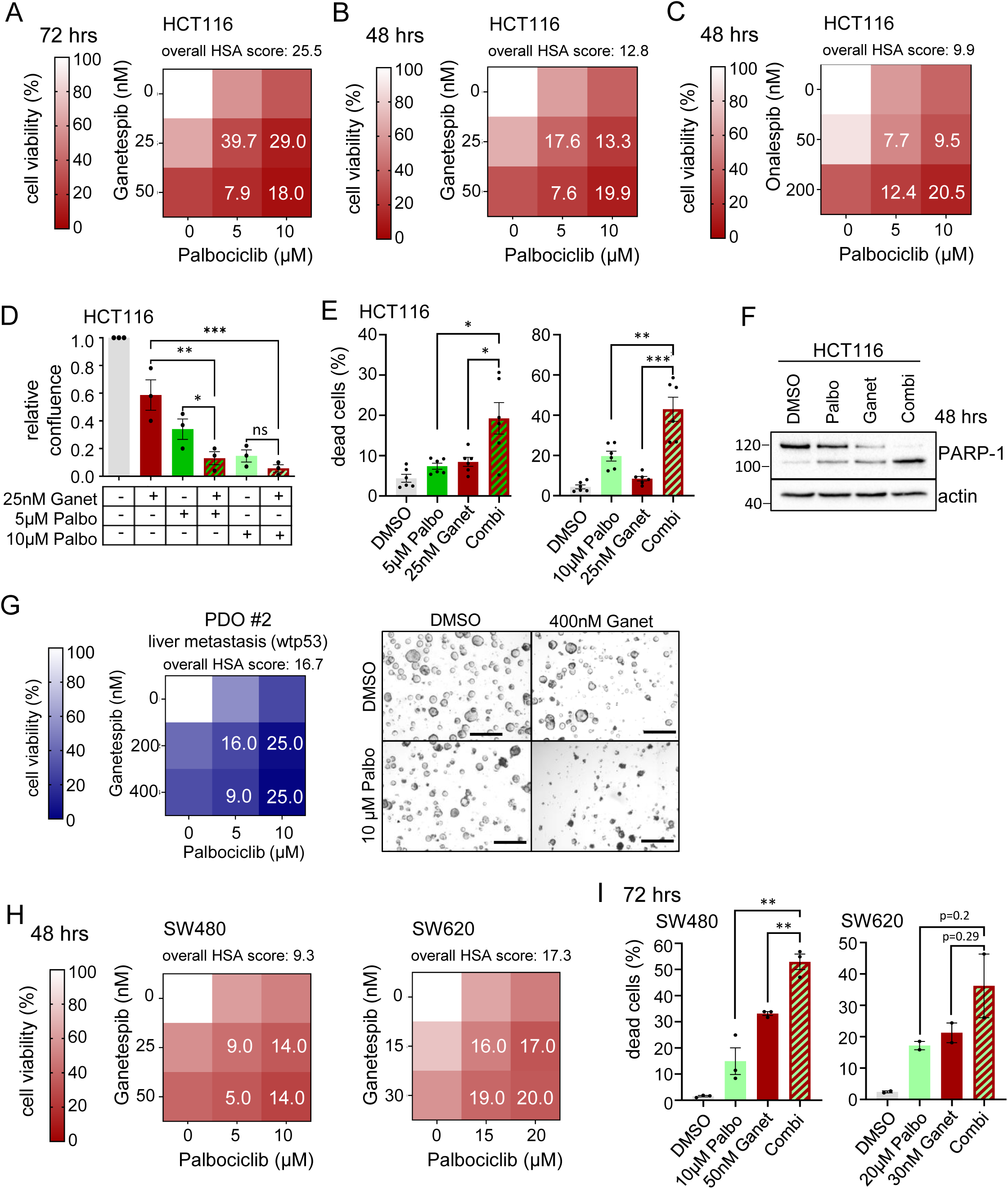
CDK4/6 inhibition in combination with HSP90 inhibitors impairs CRC cell viability independent of the p53 status. (A-C) Cell viability matrices of p53-proficient HCT116 cells treated with Palbociclib combined with Ganetespib for (A) 72 hrs or (B) 48hrs; or (C) the HSP90 inhibitor Onalespib combined with Ganetespib for 48 hrs. n ≥ 3 biological replicates. (D) Relative confluence of HCT116 cells after 72 hrs with the indicated treatments. Cell confluence was analyzed by Celigo imaging cytometry. Confluence relative to DMSO control. Palbo: Palbocliclib, Ganet: Ganetespib. (E) Determination of dead cells. PI/Hoechst/Annexin V staining of HCT116 cells treated for 72 hrs with Ganetespib and two different Palbociclib concentrations alone or in combination. Percent dead cells (PI+ only, annexin V+ only and PI+ Annexin V+ cells) were analyzed by Celigo imaging cytometry. (D, E) Mean ± SEM from ≥ 3 biological replicates. Student’s t-test, p*≤ 0.05, p**≤ 0.01, p***≤ 0.001; ns, not significant. (F) PARP-1 cleavage in HCT116 cells treated with 10 µM Palbociclib, 50 nM Ganetespib alone or in combination for 48 hrs. Representative immunoblot from 2 biological replicates. (G) Cell viability matrix of PDO #2 culture, treated as small organoids with the indicated concentrations of Ganetespib, Palbociclib alone or in combination for 72 hrs. Three replicates (different passages) with 2 in-plate technical replicates each. *Right*, Representative brightfield images after the indicated treatments. Scale bars, 200μm. (H) Cell viability matrices of p53-deficient SW480 (left) and SW620 (right) human CRC cells treated with Ganetespib - Palbociclib alone or in combination for 48 hrs. n = 3 biological replicates each. (I) Determination of dead cells as in (E) for p53-deficient SW480 and SW620 cells treated with Ganetespib and Palbociclib alone or in combination for 72 hrs. Dead cells were examined by Annexin and PI staining. Mean ± SEM from ≥ 2 biological replicates. Student’s t-test, p*≤ 0.05, p**≤ 0.01, p***≤ 0.001; ns, not significant. (A-C, G and H) Color scheme represents changes in cell viability. Numbers in the matrix are HSA synergy scores. Synergy scores: < −10 antagonistic; −10 to 10 additive, > 10 synergistic.

Since over 60% of all CRC patients harbor *TP53* mutations, we reasoned that p53-deficient CRC cells can still efficiently benefit from dual HSF1-HSP90 inhibition by direct CDK4/6 inhibition. To test this, we used SW480 and SW620 cells harboring missense p53 mutations (mutp53). SW620 cells, originally derived from a metastasis and SW480 cells, derived from the primary adenocarcinoma of the same patient, serve as model for advanced CRC tumor stage. Importantly, Palbociclib in combination with Ganetespib additively to synergistically reduced cell viability (Figs. 5H and S5K, L) and increased cell death (Fig. 5I) in all tested concentrations and time points in both p53-deficient cell lines.

Mechanistically, inhibition of CDK4/6 by Palbociclib under HSP90 inhibition again repressed HSF1 target genes (Figs. 6A and S6A, B) and lowered the levels of activated pSer326-HSF1 (Fig. 6B) in p53-proficient human CRC cells. Classical HSP90 clients were completely degraded after CDK4/6 inhibition plus Ganetespib (Fig. 6C), again demonstrating a marked HSF1-HSR suppression. Importantly, the same is true for p53-deficient human CRC cell lines (Figs. 6D-F). Palbociclib reduces HSF1 target gene expression (Figs. 6D, E) and pSer326-HSF1 levels (Fig. 6F) in combination treatments compared to HSP90 inhibition alone.

**Figure 6.**
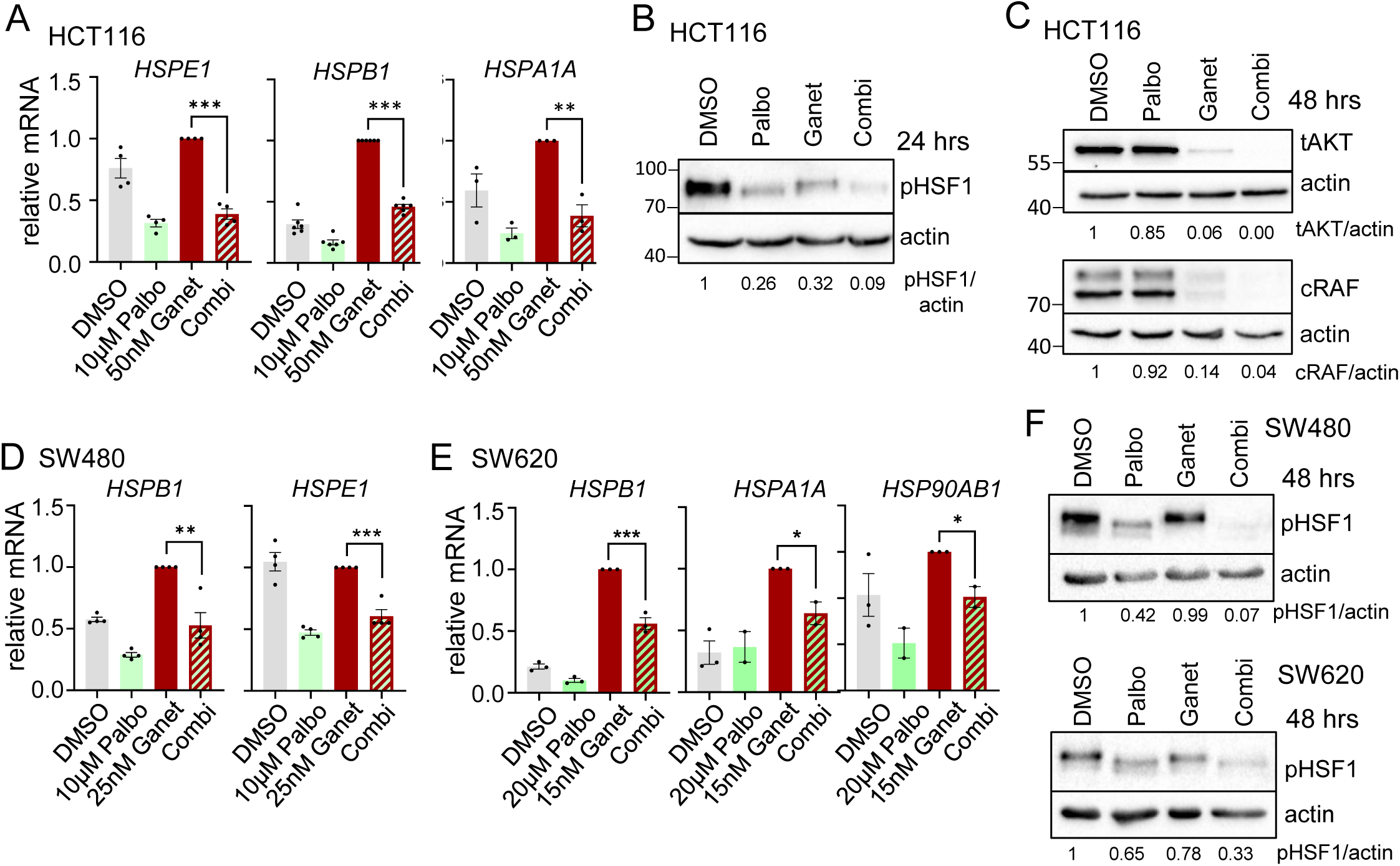
CDK4/6 inhibition in combination with HSP90 inhibitors impairs the counterproductive HSR in p53-deficient human cancer cells. (A) mRNA expression levels of representative HSF1 target genes in HCT116 cells treated as indicated for 24 hrs. qRT-PCRs, expression levels were normalized to *RPLP0* mRNA. (B, C) Immunoblots of HCT116 treated for 24 hrs (B) or for 48 hrs (C) with 50 nM Ganetespib and 10 μM Palbociclib alone or in combination. Representative of 2 biological replicates. Ganet: Ganetespib, Palbo: Palbociclib. (D, E) HSF1 target gene expression of mutp53-containing SW480 (D) and SW620 (E) cells treated with Ganetespib and Palbociclib alone or in combination for 24 hrs. qRT-PCRs, expression levels were normalized to *RPLP0* or *HPRT1* mRNA. (F) Immunoblots of p53-deficient SW480 and SW620 treated for 48 hrs with Ganetespib and Palbociclib alone or in combination. SW480 treated with 25 nM Ganet and/or 10 μM Palbo. SW620 were treated with 15 nM Ganet and/or 2.5 μM Palbo. (A, D, E) Mean ± SEM from ≥ 3 biological replicates each. Statistics relative to Ganetespib treatment. Student’s t-test, p*≤ 0.05, p**≤ 0.01, p***≤ 0.001; ns, not significant. Ganet: Ganetespib, Palbo: Palbociclib.

Next, we provided strong evidence for this strategy in p53-mutant organoids. First, p53 mutated murine organoids synergistically responded to a Palbociclib-Ganetespib combination compared to single drug treatments, including a reduction of organoid size and numbers (Fig. 7A). Second, we analyzed p53 mutant human PDOs derived from 2 different patients. Again, combined CDK4/6 inhibition by Palbociclib plus HSP90 inhibition reduced organoid viability and diminished organoid sizes compared to single drug treatments in single cell – dissociated organoids (Fig. 7B). Also, larger, multi-volume organoids responded to dual HSF1-HSP90 inhibition (Fig. 7C). Moreover, we verified the superiority of dual pathway inhibition in a matched pair of chemo-sensitive and chemo-resistant PDOs derived from another patient under CRC therapy. Even after the tumor had acquired chemo-resistance in this patient, dual CDK4/6 inhibition in combination with Ganetespib impairs organoid viability better than single treatment (Fig. 7D).

**Figure 7.**
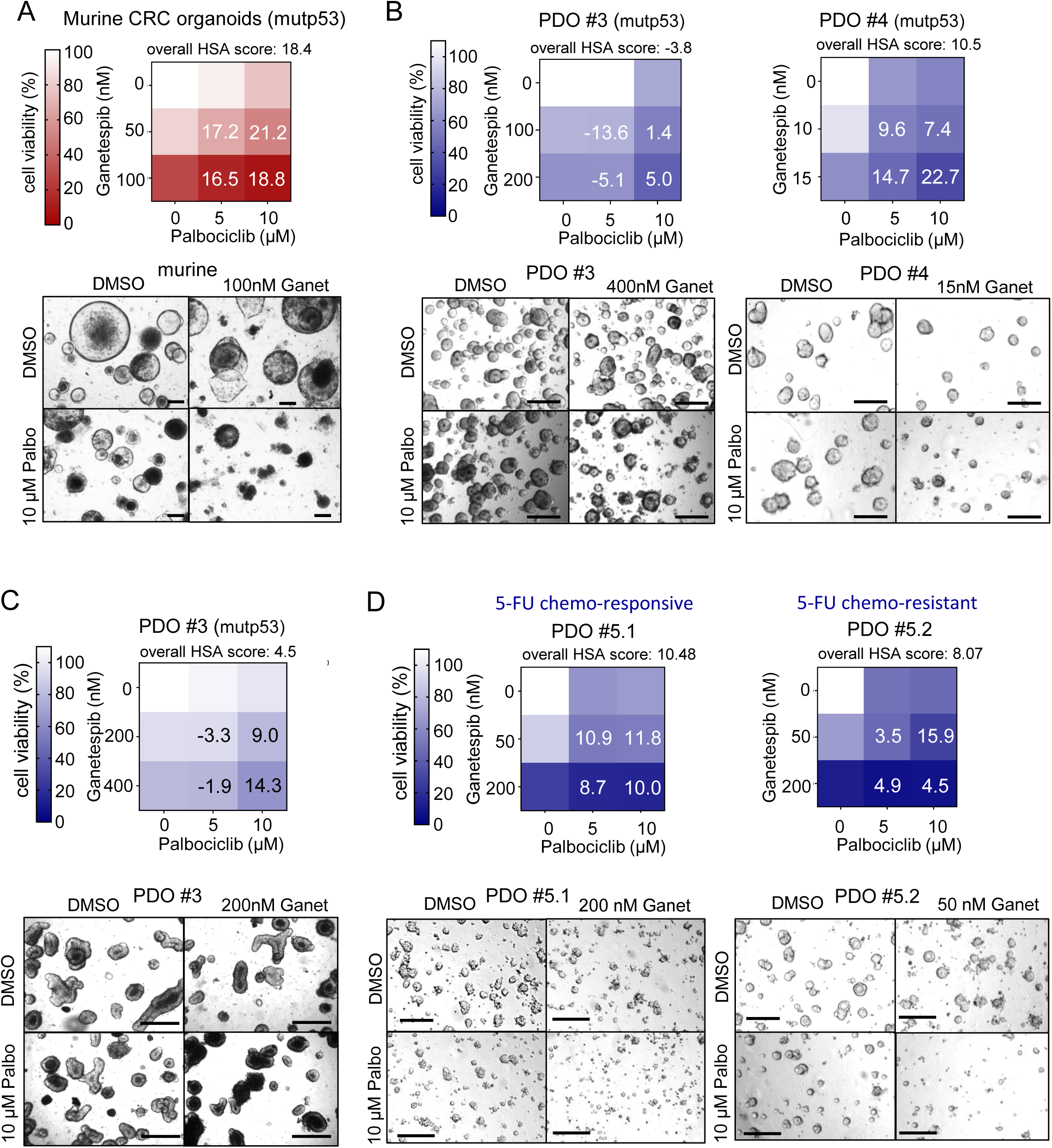
p53-mutant murine organoids and human PDOs synergistically reduce their viability after combined CDK4/6 and HSP90 inhibition. (A) Cell viability matrix of p53-mutant (mutp53) murine CRC tumor organoids treated with Ganetespib and Palbociclib alone or in combination for 72 hrs. Organoids were generated from AOM/DSS-treated *TP53*^R248Q^ mutant mice [93]. n = 3 biological replicates (different passages) with 3 in-plate technical replicates each were measured. (B) Cell viability matrices of *TP53* mutant (mutp53) patient-derived organoids (PDOs) from 2 different patients (PDO #3 and PDO #4) treated with Ganetespib, Palbociclib alone or in combination for 72 hrs (left) and 96 hrs (right). PDOs were cultivated and treated as small organoids. (C) Cell viability matrix of mutp53-containing PDO #3 treated with Ganetespib, Palbociclib alone or in combination for 48 hrs or 72 hrs. PDOs were cultivated and treated as large organoids. n = 5 biological replicates (different passages) with 3 in-plate technical replicates each were measured. (D) Cell viability matrices of a mutp53-haboring matched PDO pair (chemo-responsive and chemo-resistant) from one patient treated with Ganetespib and Palbociclib alone or in combination for 96 hrs. Both PDOs were cultivated and treated as small organoids. (A-D) Color scheme represents changes in cell viability. Numbers in the matrix are HSA synergy scores. Synergy scores: < −10 antagonistic; −10 to 10 additive; > 10 synergistic. For PDO #3 and PDO #5 cultures n ≥ 2 replicates (different passages) with ≥ 2 in-plate technical replicates each were measured. For PDO #4, one replicate was measured with 2 in-plate replicates. *Bottom*, Representative brightfield images of organoids after the indicated treatments. Scale bars, 200 μm.

In sum, these results strongly support our hypothesis that superimposed CDK4/6 inhibition prevents the unwanted HSR that contributes to therapeutic failure when treating CRC cells with HSP90 inhibitors. This work provides a strategy forward towards improving HSP90-based therapies in CRC independent of its p53 status.

## DISCUSSION

The HSF1-HSP90 system is highly upregulated and activated specifically in cancer cells but not in normal cells [8, 17, 62]. It represents a powerful pro-tumor anti-proteotoxic defense and pro-survival system, targetable with therapeutic selectivity. We and other groups have shown that HSP90 inhibition markedly impairs tumor growth and progression in preclinical mouse models *in vivo* [3–11, 16, 20, 25, 36–39]. For example, we found that HSP90 inhibition blocks progression of HER2/ErbB2-positive breast cancer [16, 25]. In the AOM/DSS CRC mouse model, HSP90 inhibition decreases tumor numbers and sizes by degradation of several HSP90 clients [20]. More advanced HSP90 inhibitors such as Ganetespib synergize with first-line therapy of ovarian cancer by interfering with DNA repair mechanisms [11]. We previously confirmed the tumor-selectivity of HSP90 inhibitors in murine colonic matched pair organoids [37]. Nevertheless, despite these encouraging findings, HSP90 inhibitors, either as monotherapy or combined with conventional chemotherapy, have so far largely failed in clinical oncology trials [6, 7, 26, 28]. One major reason for their therapeutic failure might be the abrogation of the HSP90-HSF1 negative feedback loop upon HSP90 inhibition, inadvertently leading to a deleterious rebound heat shock response and increased chaperone expression [18, 45–48]. As a result of HSP90 inhibition, tumor-driving oncoproteins might be further stabilized by alternative chaperones other than HSP90 [29–32]. Hence, there is an urgent clinical need for new strategies to block the counterproductive compensatory HSF1-mediated HSR activation upon HSP90 inhibition. Such a strategy should greatly improve the efficacy of HSP90 inhibitors. To this end, HSF1 suppression plays a key role. Notably, HSF1 is a major driver of tumorigenesis in several cancer mouse models *in vivo* [13, 63, 64]. In the AOM/DSS model of CRC, pharmacological inhibition or knock-out of HSF1 strongly suppresses colorectal carcinogenesis [64]. Moreover, in response to chronic stress, cancer cells further upregulate their global translation, thereby increasing their total proteins in an HSF1 dependent manner [65].

Building on our recent finding that HSF1 activity is effectively suppressed by wtp53 via a p53 - CDKN1A/p21 - CDK4/6 - E2F - MLK3 - MEK1 axis [33], we tested the hypothesis that p53 activation (in case of wtp53, inducing CDKN1A/p21 upregulation) or direct CDK4/6-based (in case of mutp53) cell cycle inhibition provide new options for improving the efficacy of HSP90-based therapy of CRC. Indeed, this is what we observed. Upon HSP90 inhibition the rebound HSF1 - was profoundly impaired by concomitant p53 activation in p53-proficient cancer cells or by direct HSR response CDK4/6 inhibition in p53-deficient cancer cells (Figs. 2, 3, 5, 6 and 7). Importantly, while dual HSF1-HSP90 suppression exhibits marked anti-tumoral efficacy (Figs. 1, 3, 4 and 5), normal tissues and mice showed no significant toxicities (Figs. 3C and S4B). Of note, low concentrations of HSP90 inhibitors (compared to other studies) in dual HSF1-HSP90 pathway inhibition were sufficient to strongly impair cell growth in colon-derived tumor organoids and in mice. In contrast, the same low dose as monotherapy was insufficient to adequately repress tumor cell growth.

Up to now, HSP90 inhibitors have been widely tested in combination with chemotherapies as a potential path to avoid chemoresistance, with the focus on HSP90 clients rather than on p53 status and HSR activity [38, 41, 66–72]. In contrast, the concept whether direct non-genotoxic p53 activation in combination with HSP90 inhibition decreases the deleterious HSR in CRC and suppresses cell survival had not previously been tested [66]. One of our previous studies tested first-generation HSP90 inhibitor 17-AAG plus Nutlin-3a (as p53 activator), and found that this combination destabilizes MDMX, an HSP90 client and MDM2-related p53 antagonist, causing inhibition of oncogenic survival [73]. The identified mechanism primarily caused HSP90 client degradation leading to hyperactivation of the p53 transcriptional program. However, this study did not evaluate HSF1 suppression. Based on our current work here we speculate that HSF1 target genes and HSR were also repressed, which then led to degradation of HSP90 clients such as MDMX. We found that upon dual HSF1-HSP90 pathway inhibition by p53 activation or CDK4/6 inhibition, classic HSP90 clients were even more robustly degraded than upon HSP90 inhibition alone (Figs. 2E and 6C). In support, a recent study found that concomitant CDK4/6 and HSP90 inhibition robustly destabilizes HIF1α (an HSP90 client) and decreases cell viability [74].

Interestingly, dual HSF1-HSP90 inhibition strongly activates inflammatory pathways in tumor epithelial cells (Figs. 4E, F). IL6 – JAK – STAT3, TNFα and Interferon signaling pathways were exclusively upregulated in response to drug combination, or further increased in combination treatment versus single treatment. After drug exposure in mice, we detected in the remaining tumor tissue stroma a strong decrease of cells with neutrophil characteristics, whereas other immune cells remain unchanged. Since this decrease in intratumoral neutrophils was associated with strong reductions of tumor mass in combination treatments, it can be assumed that this neutrophil population had predominantly immunosuppressive functions. Intratumoral neutrophils have been increasingly associated with protumorigenic functions [75] including angiogenesis [76] and promotion of metastasis [77]. Thus, a high ratio of neutrophils is frequently correlated with poor survival [53, 54, 78]. The observable loss of neutrophils in the course of our combination therapy, as documented by the reduced neutrophil-lymphocyte-ratio (NLR), points to a successful conversion towards a less myeloid immune phenotype consistent with intratumoral immune activation. Tumors employ several strategies to escape immune control. Based on our data, we speculate that dual HSF1-HSP90 suppression might target immune suppressive pathways by degrading immunosuppressive enzymes and/or receptors/ligands. Notably, no genes regulating immune suppression are known among HSF1 target genes. A broader analysis would be necessary to identify tumor-epithelial-derived players which contribute to the remodeling of the immune system upon dual HSF1-HSP90 inhibition.

CDK4/6 inhibition phenocopies p53 activation in p53-proficient CRC cells (Figs. 5A-F). Perhaps even more importantly, CDK4/6 inhibition also serves as alternative dual HSF1-HSP90 inhibition strategy in p53-deficient cancer cells (Figs. 5, 6 and 7), providing a strategy for improving HSP90-based tumor therapies independent of p53 status. This strategy was successful in several cases of p53-deficient patient-derived organoids (PDOs) which strongly responded to CDK4/6 inhibition under HSP90 inhibitors. Moreover, a matched PDO pair from a mutant p53 CRC patient, chemo-sensitive and chemo-resistant upon CRC therapy, impressively showed that dual HSF1-HSP90 inhibition impairs CRC survival independent of therapy status (Fig. 7D).

We propose that patients with p53-proficient tumors might be treated with a non-genotoxic p53 activator rather than a CDK4/6 inhibitor for the extra benefit of hyperactivating the powerful tumor-suppressive p53 programs (Fig. 2F). On the other hand, during long-term treatments such a combination might exert selective pressure on the tumor to acquire *TP53* mutations and favor resistance [79–81]. Proper patient monitoring should therefore be performed. If necessary, a subsequent switch to CDK4/6 inhibitors might be a second-line treatment to overcome acquired resistance mediated by *TP53* mutations in HSP90-based therapies. A further benefit of CDK4/6 inhibition is the protection of normal cells under chemotherapy, a strategy known as cyclo-therapy [82–85]. Thus, a CDK4/6-associated HSF1-HSP90 pathway inhibition in combination with chemotherapy might be useful and warrants further investigations.

Since we showed that direct p53 activation or CDK4/6 inhibition suppresses HSF1 via the E2F – MLK3 – MEK1 axis [33], we hypothesize that MAP kinase inhibition might also provide a therapeutic option in HSP90-based cancers therapies. MEK1 is a well-known upstream regulator of HSF1 [86, 87]. MEK1 inhibition in HSP90-based therapy was tested in some cancer entities yielding additive to synergistic effects in, for example, hepatocellular carcinoma (HCC) [63], non-small cell lung cancer [88] and melanoma [89], although a possible HSR abrogation again was not studied. Surprisingly, this combination has not yet been tested for CRC. As a caveat, MAP kinases often display a large degree of redundancy and cross-signaling and some, e.g. AKT and p38 kinases, also regulate HSF1 activity [90–92]. Nevertheless, in principle systematic drug screens outside the E2F – MLK3 – MEK1 axis could identify whether inhibitors of additional signaling kinases can also suppress HSF1 activity in HSP90-based therapy in CRC.

In conclusion, a HSF1 suppression by direct p53 activation or CDK4/6 inhibition improves HSP90-based therapies in CRC independent of the p53 status and in therapy-resistant CRC.

## Competing interests

The authors declare to have no competing interests.

## Funding

1. R. S.-H. is supported by the Deutsche Forschungsgemeinschaft (DFG) (SCHUH-3160/3-1). U.M.M. is supported by NCI (2R01CA176647) and the Stony Brook University TRO program. B. P. is supported by the Einstein Foundation (EVF-2018-431-2). B. P. ans S. S. are funded by the Deutsches Konsortium für Translationale Krebsforschung (DKTK) (BE01-SOB-FON-PAP).

## Author contributions

Conceptualization R. S.-H. and T. I.; Methodology T. I., K.-L. S., B. P., S. S., L.-C. C., T. De O., F. K., B. M., V. V., F. F., and R. S.-H.; Acquisition of data: T. I., K.-L. S., F. W., B. M., F. K., and R. S.-H.; Analysis and interpretation of data: all authors; Writing Original Draft T. I. and R.S-H.; Editing manuscript: R. S.-H. and U. M. M.; Reviewing & Final approval: all authors; Funding Acquisition R. S.-H.; Supervision R. S.-H.

## Acknowledgement

We thank Juline Nehring (UMG Promotionskolleg) for technical assistance.

## Ethics Statement

Patient tissue for organoid generation were provided by the Department of General, Visceral and Pediatric Surgery of the University Medical Center Göttingen (UMG, Germany) with approval from the ethics committee of UMG for the collection of CRC samples (Approval # 9/8/08 and # 25/3/17) and by the Charité – Universitätsmedizin Berlin with Approvals # EA1/161/21 and # EA1/069/113.

## METHODS

### Cell culture and treatment

RKO, HCT116+/+, HCT116-/- and SW480 cells were cultured in RPMI 1640 supplemented with 10 % fetal bovine serum, glutamine and penicillin/streptomycin. SW620 cells were cultured in Leibovitz medium supplemented with 10 % fetal bovine serum and penicillin/streptomycin. All cells besides SW620 were grown in a humidified atmosphere at 37°C with 5 % CO_2_. SW620 cells were grown under 0 % CO_2_. RG-7388 (Merck), Palbociclib (Sigma) and Ganetespib (Syntha Pharmaceuticals) were dissolved according to manufacturer’s guidelines and used as indicated.

### Mouse experiments

Animal experiments were approved by the Göttingen University Medical Center Ethik Kommission and by the Niedersächsisches Landesamt für Verbraucherschutz und Lebensmittelsicherheit, LAVES, Lower Saxony, Germany.

10-week-old male C57BL/6J mice weighing at least 20 g were used for experiments. The experiments were performed under pathogen-free barrier conditions. To induce colorectal carcinoma (CRC) a single dose of Azoxymethane (AOM, 10 mg/kg in 0.9 % sodium chloride, Sigma) was injected intraperitoneally. One-week post AOM injection, acute colitis was induced through the addition of 2 % dextran sodium sulfate (DSS, MP Biomedicals) to the drinking water for 6 days. For generating mutp53-harboring murine organoids, mice containing two humanized *TP53*^R248Q^ alleles [33, 93] were treated with 10 mg/kg AOM and 1.5% DSS for 6 days to induce CRC tumors.

5 weeks after AOM induction, tumor growth was evaluated using colonoscopy (Karl Storz GmbH). The Becker & Neurath score was used for tumor size scoring [94]: S1 = just detectable, S2 = 1/8 of the lumen, S3 = 1/4 of the lumen, S4 = 1/2 of the lumen and S5 > 1/2 of the lumen. Once mice had developed at least one S2 tumor and three S1 tumors, treatment commenced. Mice received 50 mg/kg RG-7388 (MedChem Tronica) orally (dissolved in 10 % DMSO in 0.5 % HMPC/1 % Tween 80) 5x per week, 50 mg/kg Ganetespib (Synta Pharmaceuticals) intravenously (dissolved in 10 % DMSO/18 % Cremophor RH40/3.6 % Dextrose in H_2_O) 2x per week, or their combination for the duration of 3 weeks. Using weekly colonoscopies, tumor growth was visualized to follow differences in the treatment.

After 3 weeks of treatment all mice were euthanized. Colons were harvested, opened longitudinally, and cleaned. Using a caliper, tumor sizes were measured, and tumor numbers were recorded. Where possible, tumor biopsies were taken. Colons were rolled up into “swiss rolls” and fixed in 4 % paraformaldehyde/PBS and bisected. For histological processing, both halves were placed into a single cassette and embedded in paraffin.

### Preparation and cultivation of murine small intestinal and colonic tumor organoids

Small intestinal and colonic tumor organoids were prepared as described by Klemke *et al.* [95]. In brief, 10-week old C57BL/6J mice were subjected to AOM/DSS induction of colonic tumors as described above. Once mice had developed tumors, organoids were generated from the normal small intestinal epithelium and the colon tumors. To this end the small intestinal tissue was incubated in EDTA/PBS; colonic tumor tissue was incubated in collagenase. Crypts were cultivated in Matrigel and appropriate medium at 37°C with 5 % CO_2_. Small intestinal organoid medium: advanced DMEM F-12, supplemented with 10 % Rspondin-1 conditioned medium, 20 % Noggin conditioned medium, 50 ng/mL rmEGF, 80 µM N-Acetyl-L-Cysteine, N2, B-27, HEPES, Penicillin/Streptomycin and GlutaMAX. Colonic organoid medium: advanced DMEM F-12, supplemented with 50 % Wnt3a conditioned medium, 10 % Rspondin-1 conditioned medium, 20 % Noggin conditioned medium, 200 ng/mL rmEGF, 80 µM N-Acetyl-L-Cysteine, 10 mM Nicotinamide, 500 nM A83-01, 5 µM CHIR 99021, 3.4 µg/mL ROCK inhibitor, N2, B-27, HEPES, Penicillin/Streptomycin and GlutaMAX.

### Cultivation of patient-derived colonic tumor organoids

Clinical samples for preparation of Patient-derived organoids (PDO #1 - PDO #3) were generated by the Department of General, Visceral and Pediatric Surgery of the University Medical Center Göttingen (UMG, Germany) with Approval # 9/8/08 and # 25/3/17 from the UMG Ethics Committee. PDOs were cultivated in advanced DMEM F-12, supplemented with GlutaMAX, HEPES, Penicillin/Streptomycin, 50 % Wnt3a conditioned medium, 20 % Rspondin-1 conditioned medium, 10 % Noggin conditioned medium, B27, N2, 1.25 mM N-Acetyl-L-Cysteine, 500 nM A-83-01, 10 µM SB202190, 50 ng/mL hEGF, 1 mM nicotinamide and primocin. During treatment, SB202190 and primocin were removed from the organoid medium.

For PDO #4, clinical sample was obtained at the Charité Universitätsmedizin Berlin, with Approval EA1/069/113. The matched pair of PDO #5 was obtained by the Department of Hematology, Oncology and Cancer Immunology (CCM) at the Charité Universitätsmedizin Berlin (Approval # EA1/161/21). PDOs were cultivated in advanced DMEM F-12, supplemented with GlutaMAX, HEPES, Penicillin/Streptomycin, B27, N2, 1.25 mM N-Acetyl-L-Cysteine, 500 nM A-83-01, 3 µM SB202190, 50 ng/mL hEGF, 20 ng/mL hFGF, 10 mM nicotinamide and primocin.

Large, multi-volume PDO cultures were achieved by using only mechanical disruption during the passaging procedure. Single cell dissociated, small organoid cultures were subjected to stronger dissociation during passaging. To this end organoids were incubated in Trypsin/Rho-kinase Inhibitor/DNase solution and incubated for 15 min, followed by mechanical disruption.

### Histological analysis

Murine formalin-fixed paraffin-embedded (FFPE) tissue was used for Hematoxylin and Eosin (H&E) staining by standard protocol. Tumor areas were determined using H&E stained sections and ImageJ software.

### Multiplex Immunohistochemistry (mIHC)

mIHC staining was performed according to the manufacturer’s instructions (Opal^TM^ 4-Color Anti-Rabbit Automation IHC; Opal^TM^ Polymer Anti-Rabbit HRP Secondary Antibody Kit; Akoya Biosciences^®^). The following primary rabbit antibodies were used for staining: CD4 (CST25229; 1:200), CD8 (CST98941; 1:400), F4/80 (CST 70076; 1:1000), and Ly6G (CST 87048; 1:1000) were purchased from Cell Signaling Technology^®^; CD11b (Ab133357; 1:3000) from Abcam^®^; PanCytokeratin (NBP3-07280; 1:1000) from Novus Biologicals^®^. Nuclei were stained with DAPI (Akoya Biosciences^®^). Slides were scanned using the “Vectra Polaris” platform (Akoya Biosciences^®^). Spectral unmixing and further tissue /cell segmentation as well as phenotyping were performed using the inForm Advanced Image Analysis software (inForm v2.4.10, Akoya Biosciences^®^). Multiplex data analyses: Data output from inForm 2.4.10 was further processed in RStudio (Rstudio v1.1.456) using “R” packages “Phenoptr” and “PhenoptrReports” with R version 4.1.0. Consolidated results for “Cell densities” were exported and used for further analyses.

### Confluence and Cell viability in human cancer cells

Cells were seeded in 96-well plates (Corning) and treated as described. The confluence of living cells was measured daily using the Celigo Imaging Cytometer and the Nexcelom Software v5.0.0.0.

Using CellTiter-Glo^®^ Luminescent Cell Viability Assay (Promega), cell viability was assessed according to the manufacture’s’ guidelines. Biological replicates were measured in triplicates and viability was normalized to DMSO control. HAS (Highest Single Agent) Synergy scores were calculated using the synergyfinder.fimm.fi (version 3.0) web application. Synergy scores: < −10 is antagonistic, −10 to 10 is additive, and > 10 is synergistic.

The definition of the HSA score (or Gaddum’s non-interaction model) is: “It assumes that the expected combination effect equals to the higher individual drug effect at the dose in the combination, representing the idea that a synergistic drug combination should produce additional benefits on top of what its components can achieve alone” [96].

### Cell death assay in human cancer cells

Propidium iodide (PI), Hoechst 33342 (Hoechst) and FITC Annexin V staining was performed for quantification of cell death. Cells were seeded in black clear-bottom 96-well plates. Following experimental treatments at end point, 7 µg/mL PI, 70 µg/mL Hoechst and 0.6 μg/ mL Annexin V prediluted in Annexin V binding buffer were added to the medium. Following incubation for 20 min, the Celigo Imaging Cytometer (Nexcelom) was used for the acquisition and analysis of fluorescence images. Definition for staining: ‘dead cells’ are all cells stained with Annexin V and/or PI; only Annexin V indicates early apoptosis; Annexin V + PI indicates late apoptosis; only PI indicates cell death other than apoptosis.

### Organoid morphology and organoid viability

Murine organoids and PDOs were seeded in 96-well plates (Corning) as at least two in-plate replicates and treated as described. Brightfield images were taken by the Celigo Imaging Cytometer (Nexcelom). For quantification of dead organoids, brightfield images of treated organoids were used. Dead tumor colonic organoids and small intestinal organoids were counted by visual inspection. A dead organoid was classified as one with no intact outer membrane and dark, extruded cells in the organoid lumen.

Organoid viability was measured with CellTiter-Glo^®^ 3D assay (Promega) accordingly to the manufacturer’s guidelines. Synergy scores were calculated using the synergyfinder.fimm.fi (version 3.0) or synergyfinder.org web applications. Synergy scores: < −10 is antagonistic, −10 to 10 is additive, and > 10 is synergistic.

### Immunoblots

RIPA buffer (1 % TritonX-100, 1 % Desoxycholate, 0.1 % SDS, 150 mM NaCl, 10 mM EDTA, 20 mM Tris-HCl pH7.5 and complete protease inhibitor mix, Roche) was used to prepare protein lysates. Lysates were sonicated and centrifuged, followed by BCA protein assay (Pierce) to determine protein concentrations. For SDS-polyacrylamide gel electrophoresis, equal amounts of protein were run and transferred onto nitrocellulose membranes (Millipore). The membranes were blocked with 5 % milk and incubated with the following antibodies: PARP-1, AKT and cRAF (Cell Signaling), phospho-Ser326 HSF1 and beta-Actin (Abcam). More details on antibody dilutions in Table 1. Densitometric measurements for quantification of immunoblot bands were done using the gel analysis software Image Lab™ (BioRad) and normalized to loading controls.

**Table 1:**
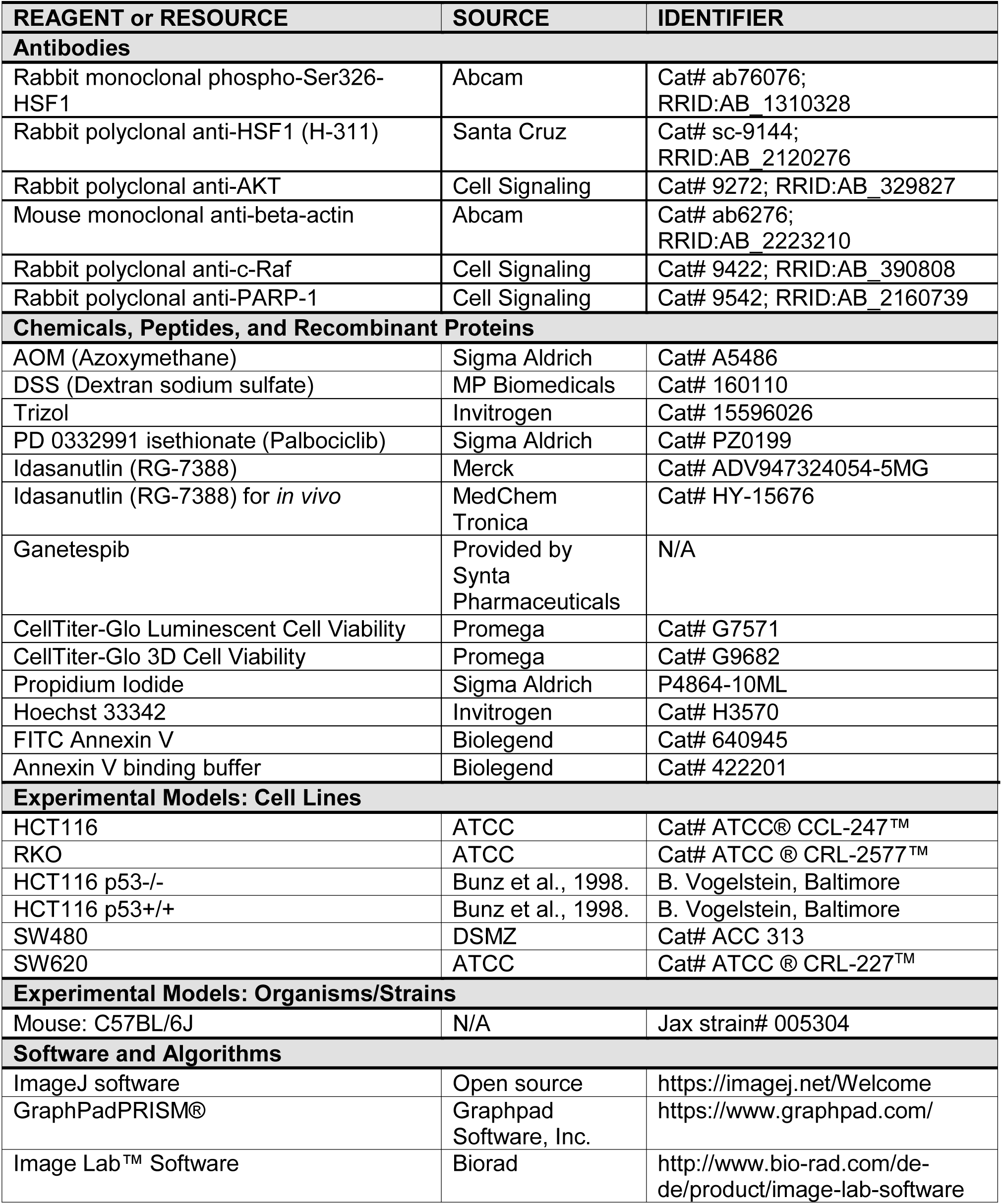
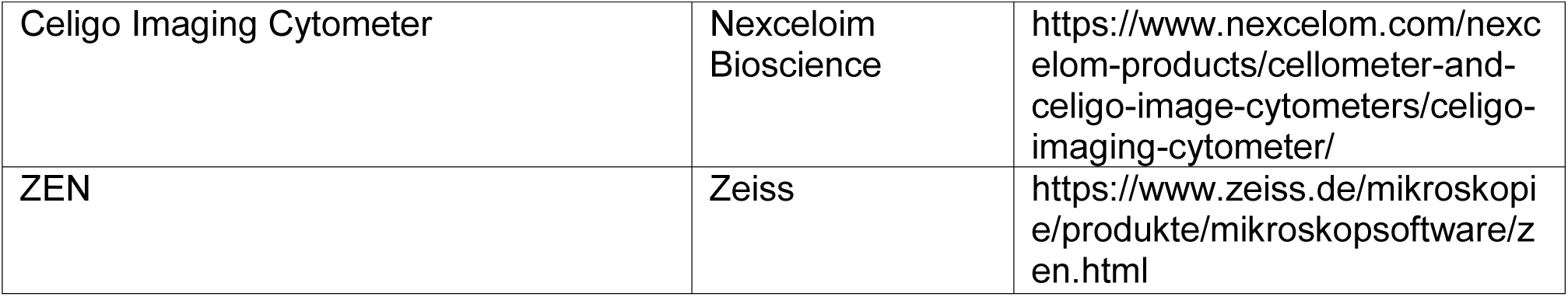
Reagents and Resources.

### Quantitative real-time PCR (qRT-PCR)

RNA from cells or organoids was isolated with Trizol according to the manufacturer’s guidelines (Invitrogen/Thermo Fisher Scientific). Equal amounts of RNA were transcribed with reverse-transcription (M-MuLV Reverse Transcriptase, NEB). Quantitative real-time PCR (qRT-PCR) analysis was performed using a SYBR green-based qPCR Master-Mix (75 mM Tris-HCl pH 8.8, 20 mM (NH_4_)_2_SO_4_, 0.01 % Tween-20, 3 mM MgCl_2_, SYBR Green 1 : 80,000, 0.2 mM dNTPs, 20 U/ml Taq-polymerase, 0.25 % TritonX-100, 300 mM Trehalose). CT values of the genes of interest were normalized to *RPLP0* mRNA. Mean ± SEM of 2 or more independent experiments, pipetted at least in duplicates. Primers are specified in Table 2.

**Table 2:**
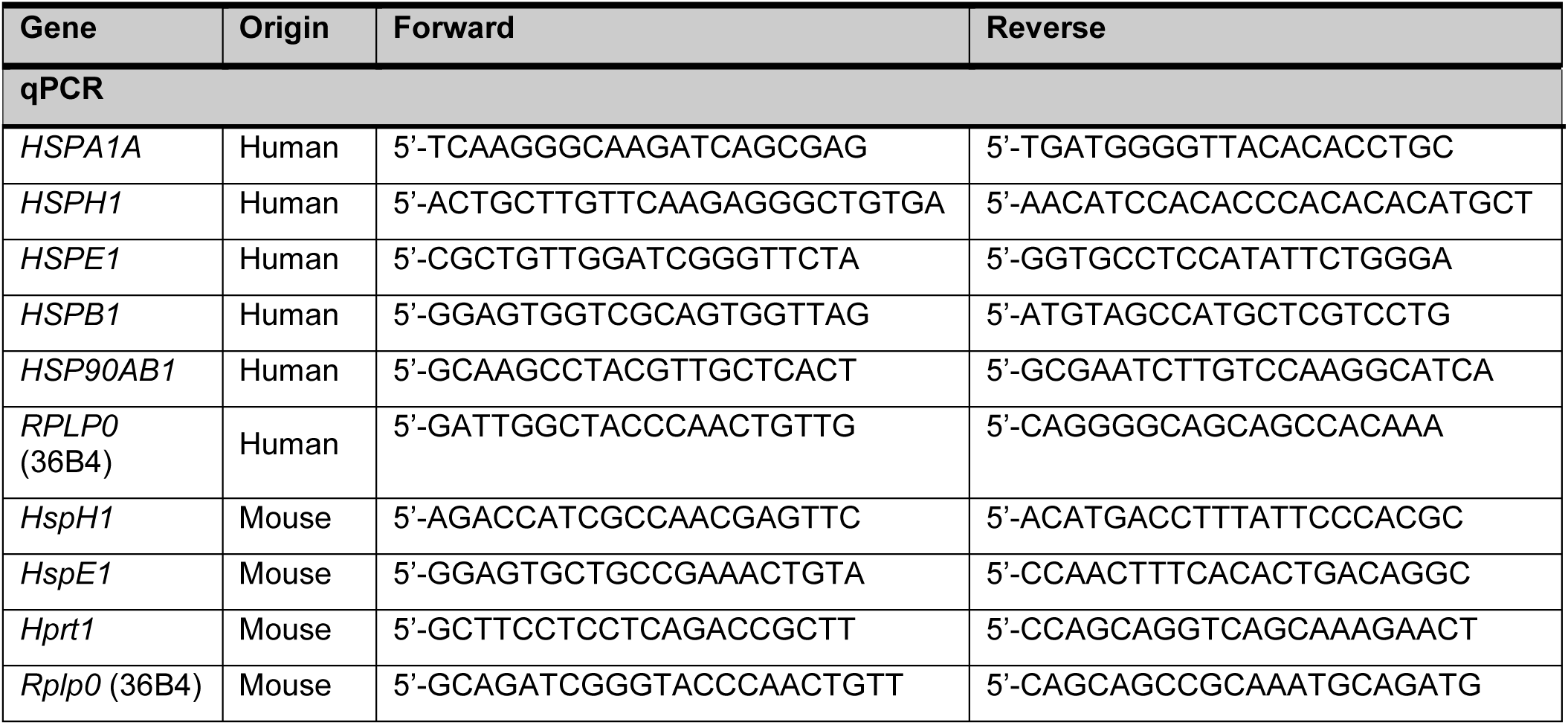
Primers for qPCR.

### mRNA sequencing of HCT116 cells

RNA samples from HCT116 treated cells were generated as pools. Two biological replicates, each consisting of two pooled biological replicates per treatment group (DMSO, Ganet, RG-7388, combi) were analyzed. RNA samples were sequenced by Novogene (Cambridge Science), including mRNA library preparation (poly A enrichment), NovaSeq 6000 PE150 sequencing, and raw Data quality control. Raw sequencing data were then processed in the high-performance computing cluster provided by the Gesellschaft für wissenschaftliche Datenverarbeitung mbH Göttingen (https://www.gwdg.de/). Quality check was performed with FastQC (version 0.11.4). Reads were trimmed (11 bp from the 5′ end, FASTQ Trimmer (FASTX toolkit version 0.0.14)), aligned to the human genome (GRCh38.106) and assigned to genomic features with RNA STAR (version 2.7). DESeq2 (version 2.11.40.6) was carried out in the GALAXY environment (https://galaxy.gwdg.de) with default parameters for differential expression analyses. For WTp53 target genes normalized counts generated by DESeq2 were used to perform GSEA analysis (Broad Institute, version 4.1.0) with the following parameters: number of permutations = 1000, type = gene_set, no_collapse, max size = 1000, and min size = 15. For HSF1 analysis, the HSF1 target gene set was extracted from Vilaboa *et al*. [97] (Supplementary Data, upregulated genes after heat-shocked HeLa cells) and uploaded into GSEA as GMX file.

### Quantification and statistical analysis

Statistics of each experiment (number of animals, number of tumors, biological replicates, technical replicates and precision measures (mean and ±SEM)) are provided in the figures and figure legends. For levels of significance, the following designations were used within this manuscript: p* ≤ 0.05; p** ≤ 0.01; p*** ≤ 0.001; ns, not significant.

---

**Supplementary Figure 1.**
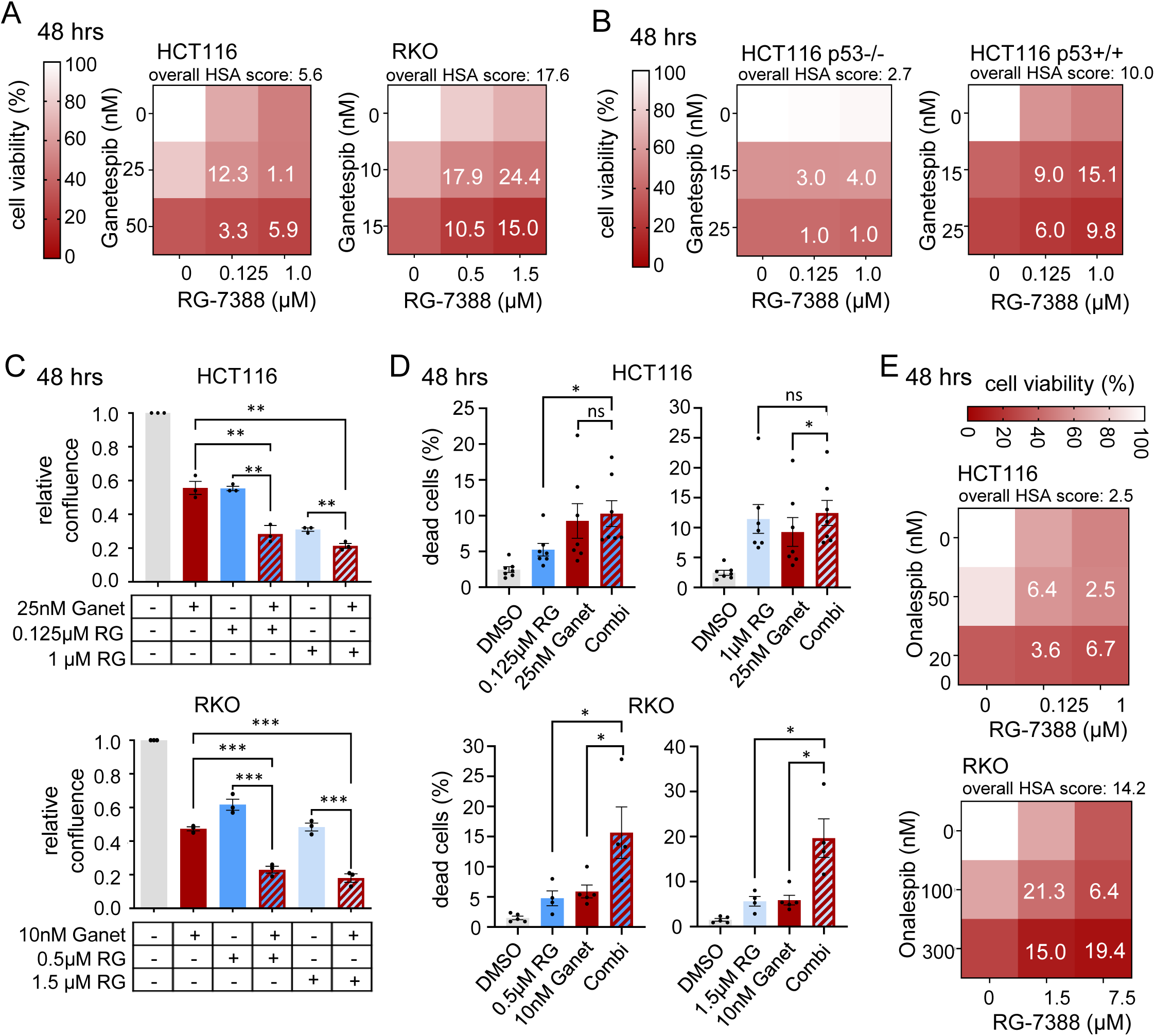
Dual HSP90-HSF1 inhibition via p53 activation synergistically impairs colorectal cancer cell growth. (A, B) Cell viability matrices of (A) HCT116 and RKO cells and (B) isogenic HCT116 p53-/- and HCT116 p53+/+ cells treated with Ganetespib – RG-7388 combinations for 48 hrs at the indicated concentrations. ≥ 3 biological replicates. (C) Relative confluence of HCT116 (top) and RKO (bottom) cells treated for 48 hrs. Cell confluence was analyzed by Celigo imaging cytometer. Confluence relative to DMSO control, set at value 1. (D) Induction of cell death. PI/Hoechst/Annexin V staining of HCT116 (top) and RKO (bottom) cells treated for 48 hrs with Ganet plus increasing concentrations of RG-7388. Percent dead cells include PI+ only, annexin V+ only and PI+ Annexin V+ cells and were analyzed using a Celigo imaging cytometer. (E) Cell viability matrices of HCT116 and RKO cells treated with Onalespib – RG-7388 combinations for 48 hrs at the indicated concentrations. ≥ 2 biological replicates. (C-D) Mean ± SEM from ≥ 3 biological replicates, Student’s t-test, p*≤ 0.05, p**≤ 0.01, p***≤ 0.001; ns, not significant. Ganet: Ganetespib, RG: RG-7388. (A, B, E) Color scheme represents changes in cell viability. Numbers within the matrix indicate the HSA synergy score. Synergy scores: < −10 antagonistic; −10 to 10 additive; > 10 synergistic.

**Supplementary Figure 2.**
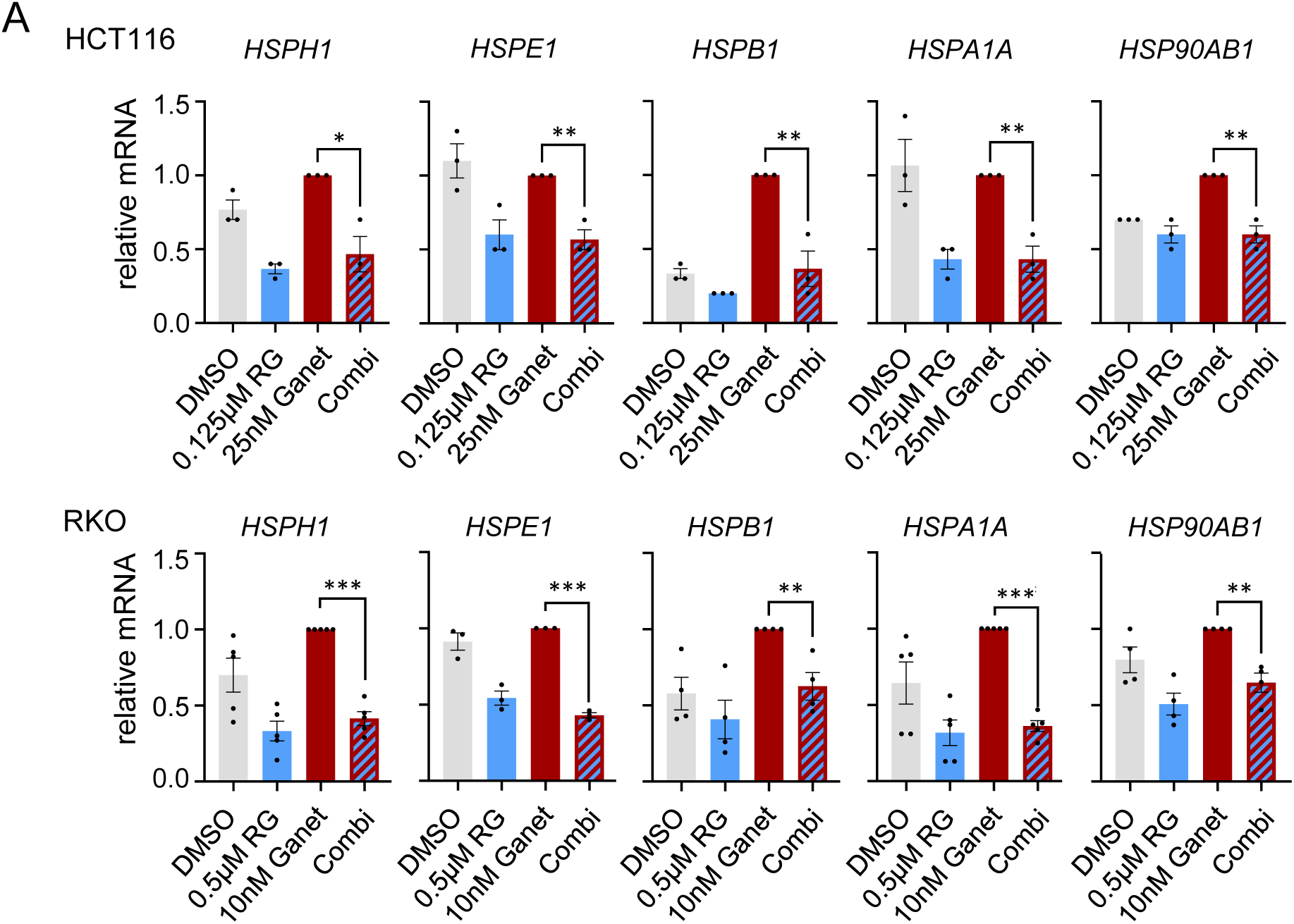
Dual HSP90-HSF1 inhibition via p53 activation abrogates the negative feedback loop. (A) mRNA expression levels of HSF1 target genes in HCT116 (top) and RKO (bottom) cells treated for 24 hrs with low concentrations of Ganetespib and RG-7388. qRT-PCRs for the indicated mRNAs normalized to *RPLP0* mRNA. Mean ± SEM from ≥ 3 biological replicates. Student’s t-test, p*≤ 0.05, p**≤ 0.01, p***≤ 0.001; ns, not significant. Ganet: Ganetespib, RG: RG-7388.

**Supplementary Figure 4.**
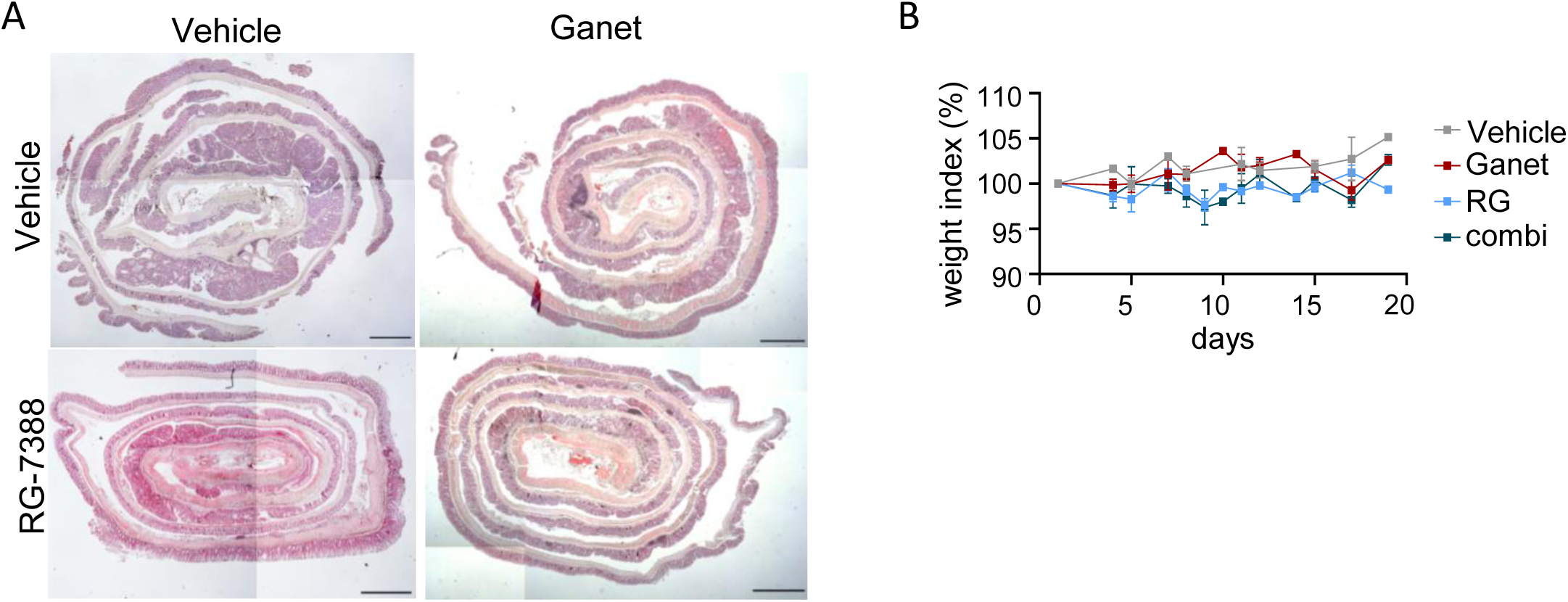
Targeting the HSF1-HSP90 pathway reduces murine CRC progression. (A) Representative images of H&E-stained cross-sections of rolled-up resected full-length colons (called swiss role) from single or combination treated C57BL6/J mice at endpoint. Mice received 50 mg/kg RG-7388 orally 5x per week, or 50 mg/kg Ganetespib intravenously 2x per week, or both. x2.5 magnification, scale bar 2 mm. (B) Lack of weight change in single or combination treated mice over the duration of the drug treatment for 19 days. Mean ± SEM from ≥ 5 mice per group. The weight of each mouse at the start of treatment was set to 100%. Y-axis starts at 90%. Ganet: Ganetespib, RG: RG-7388.

**Supplementary Figure 5.**
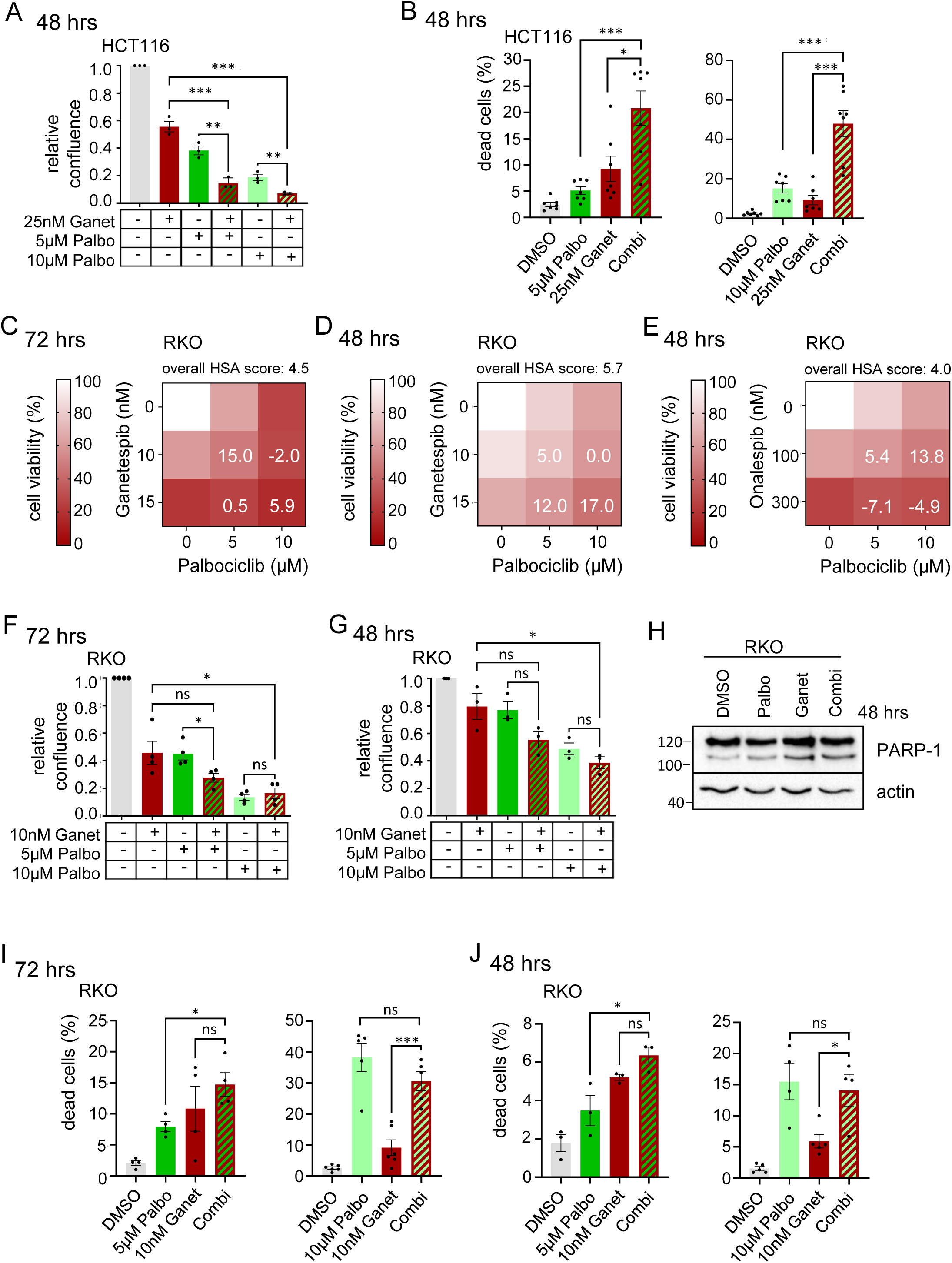
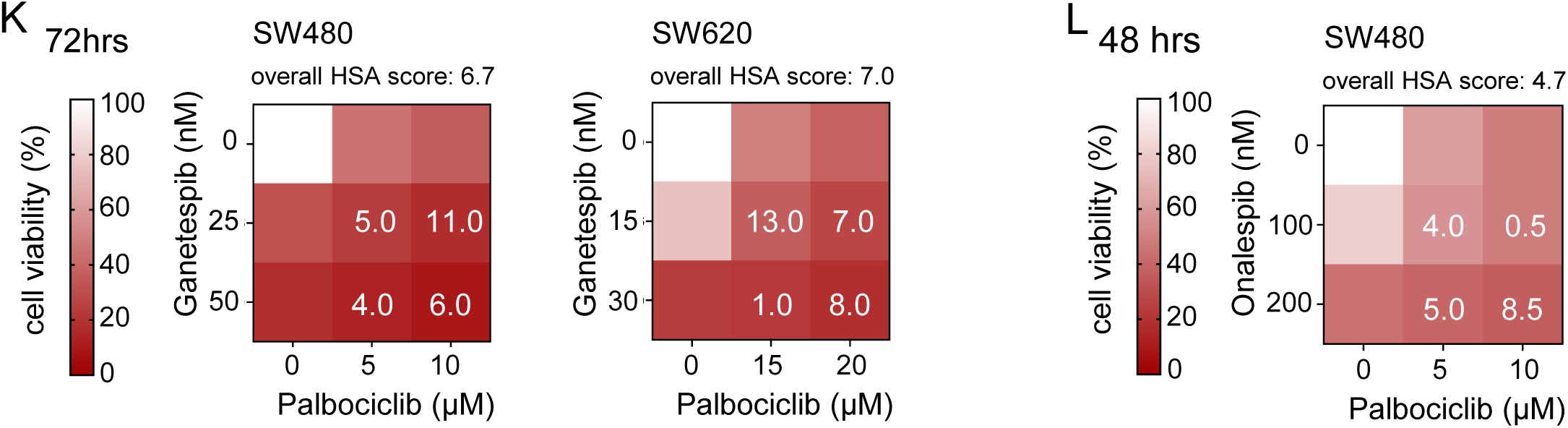
CDK4/6 inhibition in combination with HSP90 inhibitors impairs viability of CRC cancer cells independent of the p53 status. (A) Relative confluence of p53-proficient HCT116 cells treated with Ganetespib alone or in combination with Palbociclib for 48 hrs, analyzed by Celigo imaging cytometry. Confluence relative to DMSO control. Palbo: Palbocliclib, Ganet: Ganetespib. (B) Determination of dead cells treated with Ganetespib and two different Palbociclib concentrations alone or in combination for 48hrs. Percent dead cells (PI+ only, annexin V+ only and PI+ Annexin V+ cells) were analyzed using by Celigo imaging cytometry. (C-E) Cell viability matrices of RKO cells treated with Ganetespib and Palbociclib alone or in combination for (C) 72 hrs or (D) 48 hrs; or with Onalespib and Palbociclib alone or in combination for 48 hrs (E). (F, G) Relative confluence of RKO cells treated similar as in (A). (H) PARP-1 cleavage in RKO cells treated with 10 µM Palbociclib, 15 nM Ganetespib alone or in combination for 48 hrs. Representative immunoblot from 2 biological replicates. (I, J) Determination of dead cells as in (B). p53-proficient RKO cells were treated for 72 hrs (I) or 48 hrs (J) with Ganetespib and two different Palbociclib concentrations alone or in combination. Dead cells were examined by Annexin and PI staining. (K) Cell viability matrices of p53 mutant SW480 and SW620 cells treated with Ganetespib – Palbociclib alone or in combination as in (C) for 72 hrs. (L) Cell viability matrix of SW480 cells treated for 48 hrs with Onalespib and Palbociclib alone or in combination. (A, B, F, G, I, J) Mean ± SEM from ≥ 3 biological replicates. Student’s t-test, p*≤ 0.05, p**≤ 0.01, p***≤ 0.001; ns, not significant. Ganet: Ganetespib, Palbo: Palbociclib. (C, D, K) n ≥ 3 biological replicates each, (E, L) n = 2 biological replicates each. (C-E, K, L) Color scheme represents changes in cell viability. Numbers in the matrix are HSA synergy scores. Synergy scores: < −10 is antagonistic, −10 to 10 is additive, > 10 is synergistic.

**Supplementary Figure 6.**
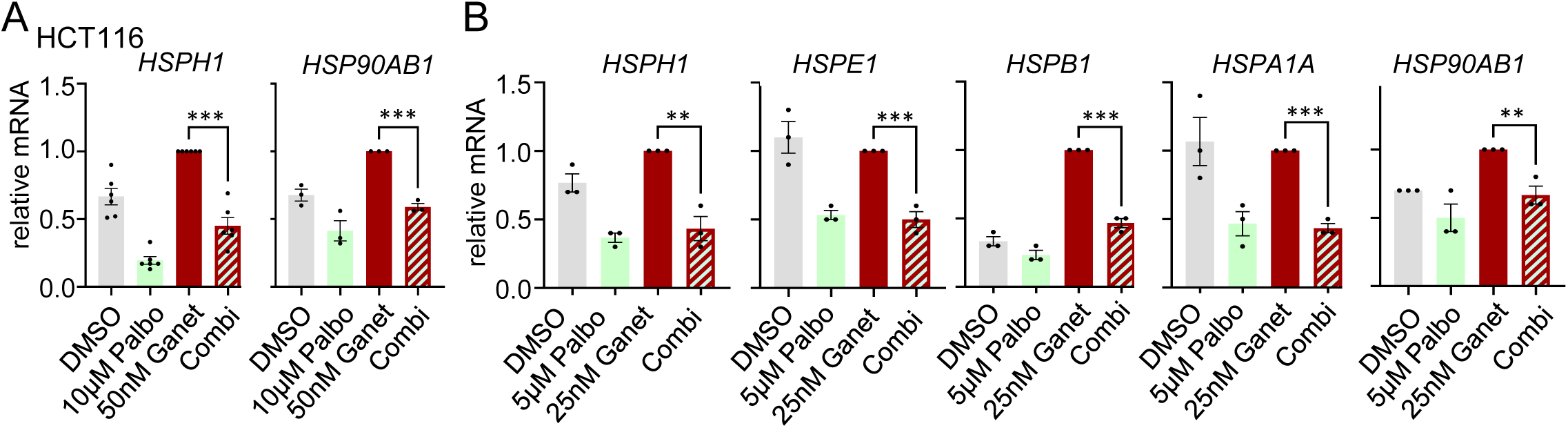
CDK4/6 inhibition in combination with HSP90 inhibitors impairs the HSR in p53-proficient cancer cells. (A, B) mRNA expression levels of representative HSF1 target genes in HCT116 cells treated for 24 hrs with the indicated concentrations of Ganetespib and Palbociclib. qRT-PCRs, expression levels normalized to *RPLP0* mRNA. Mean ± SEM from ≥ 3 biological replicates each. Student’s t-test, p*≤ 0.05, p**≤ 0.01, p***≤ 0.001; ns, not significant. Ganet: Ganetespib, Palbo: Palbociclib.

## Notes

### Competing Interest Statement

The authors have declared no competing interest.

## Literature

1. Bray, F., et al., Global cancer statistics 2018: GLOBOCAN estimates of incidence and mortality worldwide for 36 cancers in 185 countries. CA Cancer J Clin, 2018. 68(6): p. 394–424.

2. Xi, Y. and P. Xu, Global colorectal cancer burden in 2020 and projections to 2040. Transl Oncol, 2021. 14(10): p. 101174.

3. Anckar, J. and L. Sistonen, Regulation of HSF1 function in the heat stress response: implications in aging and disease. Annu Rev Biochem, 2011. 80: p. 1089–115.

4. Prince, T.L., et al., HSF1: Primary Factor in Molecular Chaperone Expression and a Major Contributor to Cancer Morbidity. Cells, 2020. 9(4).

5. Gomez-Pastor, R., E.T. Burchfiel, and D.J. Thiele, Regulation of heat shock transcription factors and their roles in physiology and disease. Nat Rev Mol Cell Biol, 2018. 19(1): p. 4–19.

6. Neckers, L., et al., Methods to validate Hsp90 inhibitor specificity, to identify off-target effects, and to rethink approaches for further clinical development. Cell Stress Chaperones, 2018. 23(4): p. 467–482.

7. Tomasic, T., et al., Discovery of Novel Hsp90 C-Terminal Inhibitors Using 3D-Pharmacophores Derived from Molecular Dynamics Simulations. Int J Mol Sci, 2020. 21(18).

8. Whitesell, L. and S.L. Lindquist, HSP90 and the chaperoning of cancer. Nat Rev Cancer, 2005. 5(10): p. 761–72.

9. Trepel, J., et al., Targeting the dynamic HSP90 complex in cancer. Nat Rev Cancer, 2010. 10(8): p. 537–49.

10. Schopf, F.H., M.M. Biebl, and J. Buchner, The HSP90 chaperone machinery. Nat Rev Mol Cell Biol, 2017. 18(6): p. 345–360.

11. Schulz-Heddergott, R. and U.M. Moll, Gain-of-Function (GOF) Mutant p53 as Actionable Therapeutic Target. Cancers (Basel), 2018. 10(6).

12. Dai, C., et al., Loss of tumor suppressor NF1 activates HSF1 to promote carcinogenesis. J Clin Invest, 2012. 122(10): p. 3742–54.

13. Dai, C., et al., Heat shock factor 1 is a powerful multifaceted modifier of carcinogenesis. Cell, 2007. 130(6): p. 1005–18.

14. Santagata, S., et al., High levels of nuclear heat-shock factor 1 (HSF1) are associated with poor prognosis in breast cancer. Proc Natl Acad Sci U S A, 2011. 108(45): p. 18378–83.

15. Hanahan, D., Hallmarks of Cancer: New Dimensions. Cancer Discov, 2022. 12(1): p. 31–46.

16. Schulz, R., et al., HER2/ErbB2 activates HSF1 and thereby controls HSP90 clients including MIF in HER2-overexpressing breast cancer. Cell Death Dis, 2014. 5: p. e980.

17. Hadizadeh Esfahani, A., et al., A systematic atlas of chaperome deregulation topologies across the human cancer landscape. PLoS Comput Biol, 2018. 14(1): p. e1005890.

18. Kijima, T., et al., HSP90 inhibitors disrupt a transient HSP90-HSF1 interaction and identify a noncanonical model of HSP90-mediated HSF1 regulation. Sci Rep, 2018. 8(1): p. 6976.

19. Li, D., et al., Functional inactivation of endogenous MDM2 and CHIP by HSP90 causes aberrant stabilization of mutant p53 in human cancer cells. Mol Cancer Res, 2011. 9(5): p. 577–88.

20. Schulz-Heddergott, R., et al., Therapeutic Ablation of Gain-of-Function Mutant p53 in Colorectal Cancer Inhibits Stat3-Mediated Tumor Growth and Invasion. Cancer Cell, 2018. 34(2): p. 298–314 e7.

21. Peng, Y., et al., Inhibition of MDM2 by hsp90 contributes to mutant p53 stabilization. J Biol Chem, 2001. 276(44): p. 40583–90.

22. King, F.W., et al., Co-chaperones Bag-1, Hop and Hsp40 regulate Hsc70 and Hsp90 interactions with wild-type or mutant p53. EMBO J, 2001. 20(22): p. 6297–305.

23. Mimnaugh, E.G., C. Chavany, and L. Neckers, Polyubiquitination and proteasomal degradation of the p185c-erbB-2 receptor protein-tyrosine kinase induced by geldanamycin. J Biol Chem, 1996. 271(37): p. 22796–801.

24. Basso, A.D., et al., Akt forms an intracellular complex with heat shock protein 90 (Hsp90) and Cdc37 and is destabilized by inhibitors of Hsp90 function. J Biol Chem, 2002. 277(42): p. 39858–66.

25. Schulz, R., et al., Inhibiting the HSP90 chaperone destabilizes macrophage migration inhibitory factor and thereby inhibits breast tumor progression. J Exp Med, 2012. 209(2): p. 275–89.

26. Shrestha, L., et al., Heat Shock Protein (HSP) Drug Discovery and Development: Targeting Heat Shock Proteins in Disease. Curr Top Med Chem, 2016. 16(25): p. 2753–64.

27. Patel, H.J., et al., Advances in the discovery and development of heat-shock protein 90 inhibitors for cancer treatment. Expert Opin Drug Discov, 2011. 6(5): p. 559–587.

28. Bickel, D. and H. Gohlke, C-terminal modulators of heat shock protein of 90kDa (HSP90): State of development and modes of action. Bioorg Med Chem, 2019. 27(21): p. 115080.

29. Kaida, A. and T. Iwakuma, Regulation of p53 and Cancer Signaling by Heat Shock Protein 40/J-Domain Protein Family Members. Int J Mol Sci, 2021. 22(24).

30. Parrales, A., et al., DNAJA1 controls the fate of misfolded mutant p53 through the mevalonate pathway. Nat Cell Biol, 2016. 18(11): p. 1233–1243.

31. Alexandrova, E.M. and U.M. Moll, Depleting stabilized GOF mutant p53 proteins by inhibiting molecular folding chaperones: a new promise in cancer therapy. Cell Death Differ, 2017. 24(1): p. 3–5.

32. Sherman, M.Y. and V.L. Gabai, Hsp70 in cancer: back to the future. Oncogene, 2015. 34(32): p. 4153–61.

33. Isermann, T., et al., Suppression of HSF1 activity by wildtype p53 creates a driving force for p53 loss-of-heterozygosity. Nat Commun, 2021. 12(1): p. 4019.

34. Kaiser, A.M. and L.D. Attardi, Deconstructing networks of p53-mediated tumor suppression in vivo. Cell Death Differ, 2018. 25(1): p. 93–103.

35. Cheok, C.F. and D.P. Lane, Exploiting the p53 Pathway for Therapy. Cold Spring Harb Perspect Med, 2017. 7(3).

36. Klemke, L., et al., The Gain-of-Function p53 R248W Mutant Promotes Migration by STAT3 Deregulation in Human Pancreatic Cancer Cells. Front Oncol, 2021. 11: p. 642603.

37. Klemke, L., et al., Hsp90-stabilized MIF supports tumor progression via macrophage recruitment and angiogenesis in colorectal cancer. Cell Death Dis, 2021. 12(2): p. 155.

38. Kramer, D., et al., Strong antitumor synergy between DNA crosslinking and HSP90 inhibition causes massive premitotic DNA fragmentation in ovarian cancer cells. Cell Death Differ, 2017. 24(2): p. 300–316.

39. Alexandrova, E.M., et al., Improving survival by exploiting tumour dependence on stabilized mutant p53 for treatment. Nature, 2015. 523(7560): p. 352–6.

40. Dobbelstein, M. and A.J. Levine, Mdm2: Open questions. Cancer Sci, 2020. 111(7): p. 2203–2211.

41. He, S., et al., The HSP90 inhibitor ganetespib has chemosensitizer and radiosensitizer activity in colorectal cancer. Invest New Drugs, 2014. 32(4): p. 577–86.

42. Desai, S., et al., Heat shock factor 1 (HSF1) controls chemoresistance and autophagy through transcriptional regulation of autophagy-related protein 7 (ATG7). J Biol Chem, 2013. 288(13): p. 9165–76.

43. Asano, Y., et al., IER5 generates a novel hypo-phosphorylated active form of HSF1 and contributes to tumorigenesis. Sci Rep, 2016. 6: p. 19174.

44. Kim, S., et al., Comparison of Cell and Organoid-Level Analysis of Patient-Derived 3D Organoids to Evaluate Tumor Cell Growth Dynamics and Drug Response. SLAS Discov, 2020. 25(7): p. 744–754.

45. Lee, J.H., et al., Heat shock protein 90 (HSP90) inhibitors activate the heat shock factor 1 (HSF1) stress response pathway and improve glucose regulation in diabetic mice. Biochem Biophys Res Commun, 2013. 430(3): p. 1109–13.

46. Samarasinghe, B., et al., Heat shock factor 1 confers resistance to Hsp90 inhibitors through p62/SQSTM1 expression and promotion of autophagic flux. Biochem Pharmacol, 2014. 87(3): p. 445–55.

47. Wang, Y. and S.R. McAlpine, N-terminal and C-terminal modulation of Hsp90 produce dissimilar phenotypes. Chem Commun (Camb), 2015. 51(8): p. 1410–3.

48. Park, S., et al., The C-terminal HSP90 inhibitor NCT-58 kills trastuzumab-resistant breast cancer stem-like cells. Cell Death Discov, 2021. 7(1): p. 354.

49. Tanaka, T., Colorectal carcinogenesis: Review of human and experimental animal studies. J Carcinog, 2009. 8: p. 5.

50. Blagih, J., et al., Cancer-Specific Loss of p53 Leads to a Modulation of Myeloid and T Cell Responses. Cell Rep, 2020. 30(2): p. 481–496 e6.

51. Guo, G., et al., Local Activation of p53 in the Tumor Microenvironment Overcomes Immune Suppression and Enhances Antitumor Immunity. Cancer Res, 2017. 77(9): p. 2292–2305.

52. Coffelt, S.B., M.D. Wellenstein, and K.E. de Visser, Neutrophils in cancer: neutral no more. Nat Rev Cancer, 2016. 16(7): p. 431–46.

53. Shen, M., et al., Tumor-associated neutrophils as a new prognostic factor in cancer: a systematic review and meta-analysis. PLoS One, 2014. 9(6): p. e98259.

54. Gentles, A.J., et al., The prognostic landscape of genes and infiltrating immune cells across human cancers. Nat Med, 2015. 21(8): p. 938–945.

55. Shaul, M.E. and Z.G. Fridlender, Tumour-associated neutrophils in patients with cancer. Nat Rev Clin Oncol, 2019. 16(10): p. 601–620.

56. Walsh, S.R., et al., Neutrophil-lymphocyte ratio as a prognostic factor in colorectal cancer. J Surg Oncol, 2005. 91(3): p. 181–4.

57. Malietzis, G., et al., The emerging role of neutrophil to lymphocyte ratio in determining colorectal cancer treatment outcomes: a systematic review and meta-analysis. Ann Surg Oncol, 2014. 21(12):p. 3938–46.

58. Zheng, W., et al., Tumor-Associated Neutrophils in Colorectal Cancer Development, Progression and Immunotherapy. Cancers (Basel), 2022. 14(19).

59. Rivlin, N., et al., Mutations in the p53 Tumor Suppressor Gene: Important Milestones at the Various Steps of Tumorigenesis. Genes Cancer, 2011. 2(4): p. 466–74.

60. Lopez, I., et al., Different mutation profiles associated to P53 accumulation in colorectal cancer. Gene, 2012. 499(1): p. 81–7.

61. Smith, G., et al., Mutations in APC, Kirsten-ras, and p53--alternative genetic pathways to colorectal cancer. Proc Natl Acad Sci U S A, 2002. 99(14): p. 9433–8.

62. Mendillo, M.L., et al., HSF1 drives a transcriptional program distinct from heat shock to support highly malignant human cancers. Cell, 2012. 150(3): p. 549–62.

63. Min, J.N., et al., Selective suppression of lymphomas by functional loss of Hsf1 in a p53-deficient mouse model for spontaneous tumors. Oncogene, 2007. 26(35): p. 5086–97.

64. Li, J., et al., Heat Shock Factor 1 Epigenetically Stimulates Glutaminase-1-Dependent mTOR Activation to Promote Colorectal Carcinogenesis. Mol Ther, 2018. 26(7): p. 1828–1839.

65. Maiti, S., et al., Hsf1 and the molecular chaperone Hsp90 support a ‘rewiring stress response’ leading to an adaptive cell size increase in chronic stress. Elife, 2023. 12.

66. Kryeziu, K., et al., Combination therapies with HSP90 inhibitors against colorectal cancer. Biochim Biophys Acta Rev Cancer, 2019. 1871(2): p. 240–247.

67. Ewers, K.M., et al., HSP90 Inhibition Synergizes with Cisplatin to Eliminate Basal-like Pancreatic Ductal Adenocarcinoma Cells. Cancers (Basel), 2021. 13(24).

68. Kim, W.Y., et al., Targeting heat shock protein 90 overrides the resistance of lung cancer cells by blocking radiation-induced stabilization of hypoxia-inducible factor-1alpha. Cancer Res, 2009. 69(4): p. 1624–32.

69. Moser, C., et al., Blocking heat shock protein-90 inhibits the invasive properties and hepatic growth of human colon cancer cells and improves the efficacy of oxaliplatin in p53-deficient colon cancer tumors in vivo. Mol Cancer Ther, 2007. 6(11): p. 2868–78.

70. Bull, E.E., et al., Enhanced tumor cell radiosensitivity and abrogation of G2 and S phase arrest by the Hsp90 inhibitor 17-(dimethylaminoethylamino)-17-demethoxygeldanamycin. Clin Cancer Res, 2004. 10(23): p. 8077–84.

71. Modi, S., et al., HSP90 inhibition is effective in breast cancer: a phase II trial of tanespimycin (17-AAG) plus trastuzumab in patients with HER2-positive metastatic breast cancer progressing on trastuzumab. Clin Cancer Res, 2011. 17(15): p. 5132–9.

72. Banerji, U., et al., Correction to: An in vitro and in vivo study of the combination of the heat shock protein inhibitor 17-allylamino-17-demethoxygeldanamycin and carboplatin in human ovarian cancer models. Cancer Chemother Pharmacol, 2018. 82(5): p. 911–912.

73. Vaseva, A.V., et al., Blockade of Hsp90 by 17AAG antagonizes MDMX and synergizes with Nutlin to induce p53-mediated apoptosis in solid tumors. Cell Death Dis, 2011. 2(5): p. e156.

74. Zhao, S., et al., Anti-cancer efficacy including Rb-deficient tumors and VHL-independent HIF1alpha proteasomal destabilization by dual targeting of CDK1 or CDK4/6 and HSP90. Sci Rep, 2021. 11(1): p. 20871.

75. Houghton, A.M., et al., Neutrophil elastase-mediated degradation of IRS-1 accelerates lung tumor growth. Nat Med, 2010. 16(2): p. 219–23.

76. Nozawa, H., C. Chiu, and D. Hanahan, Infiltrating neutrophils mediate the initial angiogenic switch in a mouse model of multistage carcinogenesis. Proc Natl Acad Sci U S A, 2006. 103(33): p. 12493–8.

77. Kowanetz, M., et al., Granulocyte-colony stimulating factor promotes lung metastasis through mobilization of Ly6G+Ly6C+ granulocytes. Proc Natl Acad Sci U S A, 2010. 107(50): p. 21248–55.

78. Templeton, A.J., et al., Prognostic role of neutrophil-to-lymphocyte ratio in solid tumors: a systematic review and meta-analysis. J Natl Cancer Inst, 2014. 106(6): p. dju124.

79. Skalniak, L., et al., Prolonged Idasanutlin (RG7388) Treatment Leads to the Generation of p53-Mutated Cells. Cancers (Basel), 2018. 10(11).

80. Aziz, M.H., H. Shen, and C.G. Maki, Acquisition of p53 mutations in response to the non-genotoxic p53 activator Nutlin-3. Oncogene, 2011. 30(46): p. 4678–86.

81. Kucab, J.E., et al., Nutlin-3a selects for cells harbouring TP53 mutations. Int J Cancer, 2017. 140(4):p. 877–887.

82. Blume, J., et al., CDK4/6 inhibition confers protection of normal gut epithelia against gemcitabine and the active metabolite of irinotecan. Cell Cycle, 2023. 22(13): p. 1563–1582.

83. Blagosklonny, M.V., Selective protection of normal cells from chemotherapy, while killing drug-resistant cancer cells. Oncotarget, 2023. 14: p. 193–206.

84. van Leeuwen, I.M., Cyclotherapy: opening a therapeutic window in cancer treatment. Oncotarget, 2012. 3(6): p. 596–600.

85. Blagosklonny, M.V. and Z. Darzynkiewicz, Cyclotherapy: protection of normal cells and unshielding of cancer cells. Cell Cycle, 2002. 1(6): p. 375–82.

86. Park, M.A., et al., Mitogen-activated protein kinase kinase 1/2 inhibitors and 17-allylamino-17-demethoxygeldanamycin synergize to kill human gastrointestinal tumor cells in vitro via suppression of c-FLIP-s levels and activation of CD95. Mol Cancer Ther, 2008. 7(9): p. 2633–48.

87. Acquaviva, J., et al., Targeting KRAS-mutant non-small cell lung cancer with the Hsp90 inhibitor ganetespib. Mol Cancer Ther, 2012. 11(12): p. 2633–43.

88. Sato, S., N. Fujita, and T. Tsuruo, Modulation of Akt kinase activity by binding to Hsp90. Proc Natl Acad Sci U S A, 2000. 97(20): p. 10832–7.

89. Lu, W.C., et al., AKT1 mediates multiple phosphorylation events that functionally promote HSF1 activation. FEBS J, 2022.

90. Dayalan Naidu, S., et al., Heat Shock Factor 1 Is a Substrate for p38 Mitogen-Activated Protein Kinases. Mol Cell Biol, 2016. 36(18): p. 2403–17.

91. Stramucci, L., A. Pranteda, and G. Bossi, Insights of Crosstalk between p53 Protein and the MKK3/MKK6/p38 MAPK Signaling Pathway in Cancer. Cancers (Basel), 2018. 10(5).

92. Tanaka, T., Development of an inflammation-associated colorectal cancer model and its application for research on carcinogenesis and chemoprevention. Int J Inflam, 2012. 2012: p. 658786.

93. Hanel, W., et al., Two hot spot mutant p53 mouse models display differential gain of function in tumorigenesis. Cell Death Differ, 2013. 20(7): p. 898–909.

94. Becker, C., M.C. Fantini, and M.F. Neurath, High resolution colonoscopy in live mice. Nat Protoc, 2006. 1(6): p. 2900–4.

95. Klemke, L., et al., Preparation and Cultivation of Colonic and Small Intestinal Murine Organoids Including Analysis of Gene Expression and Organoid Viability. Bio Protoc, 2022. 12(2): p. e4298.

96. Berenbaum, M.C., What is synergy? Pharmacol Rev, 1989. 41(2): p. 93–141.

97. Vilaboa, N., et al., New inhibitor targeting human transcription factor HSF1: effects on the heat shock response and tumor cell survival. Nucleic Acids Res, 2017. 45(10): p. 5797–5817.

